# Developmental exposure to the organochlorine pesticide dieldrin causes male-specific exacerbation of α-synuclein-preformed fibril-induced toxicity and motor deficits

**DOI:** 10.1101/2020.02.03.932681

**Authors:** Aysegul O. Gezer, Joseph Kochmanski, Sarah E. VanOeveren, Allyson Cole-Strauss, Christopher J Kemp, Joseph R. Patterson, Kathryn M. Miller, Nathan C. Kuhn, Danielle E. Herman, Alyssa McIntire, Jack W. Lipton, Kelvin C. Luk, Sheila M. Fleming, Caryl E. Sortwell, Alison I. Bernstein

## Abstract

Human and animal studies have shown that exposure to the organochlorine pesticide dieldrin is associated with increased risk of Parkinson’s disease (PD). Previous work showed that developmental dieldrin exposure increased neuronal susceptibility to MPTP toxicity in male C57BL/6 mice, possibly via changes in dopamine (DA) packaging and turnover. However, the relevance of the MPTP model to PD pathophysiology has been questioned. We therefore studied dieldrin-induced neurotoxicity in the α-synuclein (α-syn)-preformed fibril (PFF) model, which better reflects the α-syn pathology and toxicity observed in PD pathogenesis. Specifically, we used a “two-hit” model to determine whether developmental dieldrin exposure increases susceptibility to α-syn PFF-induced synucleinopathy. Dams were fed either dieldrin (0.3 mg/kg, every 3-4 days) or vehicle corn oil starting 1 month prior to breeding and continuing through weaning of pups at postnatal day 22. At 12 weeks of age, male and female offspring received intrastriatal PFF or control saline injections. Consistent with the male-specific increased susceptibility to MPTP, our results demonstrate that developmental dieldrin exposure exacerbates PFF-induced toxicity in male mice only. Specifically, in male offspring, dieldrin exacerbated PFF-induced motor deficits on the challenging beam and increased DA turnover in the striatum 6 months after PFF injection. However, male offspring showed neither exacerbation of phosphorylated α-syn (pSyn) aggregation in the substantia nigra (SN) at 1 or 2 months post-PFF injection, nor exacerbation of PFF-induced TH and NeuN loss in the SN 6 months post-PFF injection. Collectively, these data indicate that developmental dieldrin exposure produces a male-specific increase in neuronal vulnerability to synucleinopathy. This sex-specific result is consistent with both previous work in the MPTP model, our previously reported sex-specific effects of this exposure paradigm on the male and female epigenome, and the higher prevalence and more severe course of PD in males. The novel two-hit environmental toxicant/PFF exposure paradigm established in this project can be used to explore the mechanisms by which other PD-related exposures alter neuronal vulnerability to synucleinopathy in sporadic PD.

**Highlights:**

- Developmental dieldrin exposure increases α-syn-PFF-induced motor deficits
- Developmental dieldrin exposure increases PFF-induced deficits in DA handling
- Developmental dieldrin exposure does not affect PFF-induced loss of nigral neurons
- This is a novel paradigm modeling how environmental factors increase risk of PD
- Female mice show PFF-induced pathology, but no PFF-induced motor deficits.

## Introduction

Parkinson’s disease (PD), the second most common neurodegenerative disorder in the United States, is characterized by progressive degeneration of dopaminergic neurons of the nigrostriatal pathway and the formation of alpha-synuclein (α-syn)-containing Lewy bodies. Several genes have been linked to inherited forms of PD; however, it is estimated that only 5-10% of PD cases are familial.^1,2^ The remaining ∼90% of sporadic PD cases are likely due to a complex interaction between genes and environmental factors. Supporting this idea, epidemiologic studies have shown an association between exposure to persistent organic pollutants, including pesticides and industrial toxicants, and an increased risk of PD.^3–19^ When these data are combined with post-mortem analysis and mechanistic studies, a role for specific compounds in PD emerges.^12,15,20^

Dieldrin is an organochlorine pesticide that has been associated with an increased risk of PD by both epidemiologic and mechanistic studies.^15,20–25^ Because dieldrin was phased out in the 1970s and 1980s, the potential for new, acute exposure to dieldrin is low. However, the health effects of past exposures will continue for decades as the population currently diagnosed with PD and those who will develop PD in the next 20-30 years were likely exposed to dieldrin prior to its phase out.^21,26–28^ Furthermore, well-established experimental models of dieldrin exposure have demonstrated that dieldrin induces oxidative stress, is selectively toxic to dopaminergic cells, disrupts striatal dopamine (DA) activity, and may promote α-syn aggregation.^20,21,29–34^

Because of the established association of dieldrin with PD risk and well-characterized animal exposure dosing paradigms, our lab utilizes the developmental dieldrin exposure as a representative model of increased PD susceptibility.^33–35^ In this model, developmental exposure to dieldrin induces persistent alterations in the DA system that cause a male-specific increase in susceptibility to subsequent exposure to MPTP.^33^ However, numerous therapeutics that protect against MPTP in preclinical studies have failed to translate to clinical benefit, suggesting that this model has limited utility for accurately predicting clinical translation or exploring toxicological mechanisms in PD.^36^ Moreover, MPTP is a fast-acting toxicant that induces rapid and extensive loss of striatal DA, which does not reflect the protracted course of loss of function and degeneration observed in disease. Finally, the failure of the MPTP model to develop widespread α-syn pathology calls into question its validity as a “second hit” for examining organochlorine-induced PD vulnerability.^36,37^ Instead, in the present study, we incorporated the α-syn pre-formed fibril (PFF) model to investigate dieldrin-induced parkinsonian susceptibility.

In 2012, Luk et al reported that intrastriatal injection of synthetic α-syn PFFs into wild-type mice seeded endogenous accumulation of Lewy Body (LB)-like intracellular α-syn inclusions and ultimately led to nigrostriatal degeneration.^38^ These findings have been replicated in transgenic mice, non-transgenic mice, rats, and monkeys.^38–44^ PFF-induced α-syn inclusions resemble LBs in that they are compact intracytoplasmic structures of hyperphosphorylated (ser129) α-syn (psyn), co-localize with ubiquitin and p62, and are thioflavin-S-positive and proteinase-k resistant. Over time, the α-syn aggregates progressively compact and eventually lead to neuronal degeneration.^38,39,45,46^ Thus, the intrastriatal injection of α-syn PFFs can be used to model pathological synucleinopathy and nigrostriatal toxicity in mice. Here, we tested the hypothesis that developmental dieldrin exposure increases susceptibility to synucleinopathy and associated toxicity in the α-syn PFF model.

## Materials and Methods

### Animals

Male (11 weeks old) and female (7 weeks old) C57BL/6 mice were purchased from Jackson Laboratory (Bar Harbor, Maine). After a week of habituation, mice were switched to a 12:12 reverse light/dark cycle for the duration of the study. Mice were housed in Thoren ventilated caging systems with automatic water and 1/8-inch Bed-O-Cobs bedding with Enviro-Dri for enrichment. Food and water were available ad libitum. Mice were maintained on standard LabDiet 5021 chow (LabDiet). F0 females were individually housed during dieldrin dosing, except during the mating phase. F1 pups were group housed by sex; with 2-4 animals per cage. All procedures were conducted in accordance with the National Institutes of Health Guide for Care and Use of Laboratory Animals and approved by the Institutional Animal Care and Use Committee at Michigan State University.

### Dieldrin exposure paradigm

Dosing was carried out as previously described.^35^ Female mice were habituated to peanut butter feeding for three days. During this period, each mouse was fed a peanut butter pellet containing 6 µl vehicle (corn oil) and monitored to ensure peanut butter consumption. Following these three days of habituation, mice were administered 0.3 mg/kg dieldrin (ChemService) dissolved in corn oil vehicle and mixed with peanut butter pellets every 3 days.^47^ Control mice received an equivalent amount of corn oil vehicle in peanut butter. This dose was based on previous results showing low toxicity, but clear developmental effects.^33^ Consumption of peanut butter pellets was ensured via visual inspection and typically occurred within minutes. Adult C57BL/6 (8-week-old) female animals were treated throughout breeding, gestation, and lactation (Figure 1). Four weeks into female exposure, unexposed C57BL/6 males (12 weeks old) were introduced for breeding. Mating was scheduled for a maximum age difference of 2 weeks, although all females were pregnant by the end of the first week. Offspring were weaned at postnatal day 22 and separated by litter and by sex. At 12-14 weeks of age, one set of male and female offspring from independent litters was sacrificed (n=10 per treatment per sex). An additional set of animals underwent saline or PFF injections at 12 weeks of age; these animals were sacrificed at 1, 2 and 6 months post-PFF injection (n = 10 per treatment per sex). This produced 4 experimental groups (vehicle:saline, vehicle:PFF, dieldrin:saline, and dieldrin:PFF). Animals were singly housed following surgeries for the duration of the experiment.

**Figure 1:**
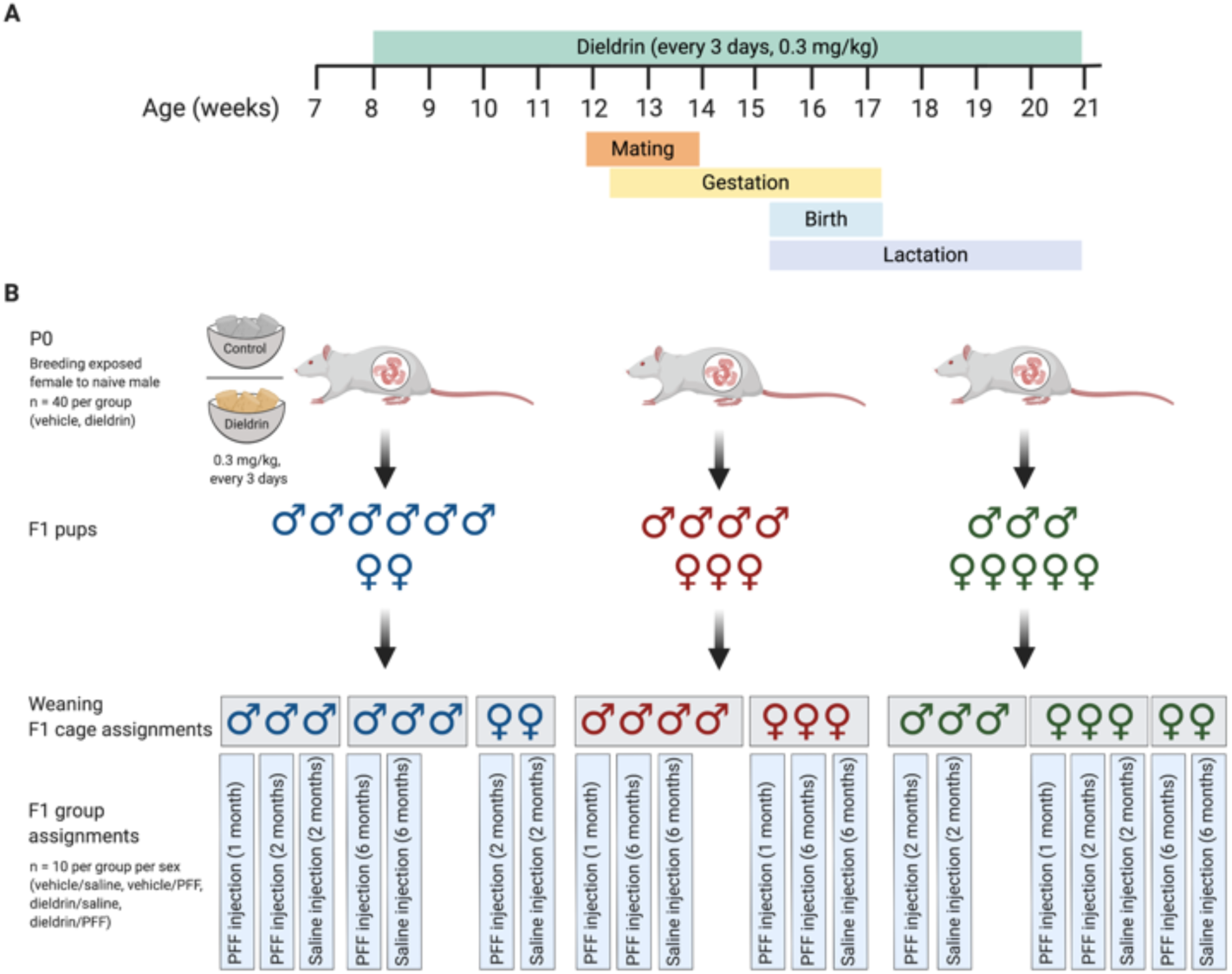
Dosing timeline, weaning strategy, cage and group assignments. A) Timeline of developmental dieldrin exposure model: In this paradigm, only female dams were fed dieldrin. Exposure began at 8 weeks of age with 0.3 mg/kg dieldrin dissolved in a corn oil vehicle and administered via peanut butter pellet. Males were introduced for mating when females were 12 weeks of age. Pregnancy was confirmed by monitoring weight. Dieldrin administration continued until pups (F1) were weaned at PND22. B) Weaning strategy, cage and group assignments for PFF injections: At weaning, pups (F1) were separated by sex and litter (colors represent different litters) with no more than 4 animals per cage (grey boxes represent cages). No animals that were singly housed were used in this study. Within each cage, animals were assigned to groups such that for every experimental group, all animals were from independent litters. Created in BioRender.

### Preparation of α-syn PFFs and verification of fibril size

Fibrils were generated using wild-type, full-length, recombinant mouse α-syn monomers, as previously described.^38,44,48–51^ Quality control was performed on full length fibrils to confirm fibril formation (by transmission electron microscopy), amyloid content (by thioflavin T assay), a shift to insoluble species compared to monomers (by sedimentation assay), and low bacterial contamination (<0.5 endotoxin units mg^-1^ of total protein via a *Limulus* amebocyte lysate assay). On the day of surgery, PFFs were thawed to room temperature, diluted to 2 μg/μl in Dulbecco’s phosphate buffered saline (PBS) (Gibco), and sonicated at room temperature using an ultrasonic homogenizer (Q125 Sonicator; Qsonica, Newtown, CT. Sonication was performed for 1 minute, using 1 second pulses with 1 second between pulses and amplitude set at 30%. Prior to surgeries, an aliquot of sonicated PFFs was analyzed using transmission electron microscopy.

### Transmission electron microscopy (TEM)

TEM was performed as described previously.^50^ Briefly, samples were prepared on Formvar/carbon-coated copper grids (EMSDIASUM, FCF300-Cu) that were washed twice by floating on drops of distilled H2O. Grids were then floated for 1 min on 10 μl drops of sonicated PFFs diluted 1:50 in PBS, followed by 1 min on 10 μl drops of aqueous 2% uranyl acetate, wicking away liquid with filter paper after each step. Grids were allowed to dry before imaging with a JEOL JEM-1400+ transmission electron microscope. Prior to intrastriatal injections of the mouse α-syn PFFs, size of the sonicated α-syn PFFs was screened using transmission electron microscopy to ensure fibril length was approximately 50 nm, a length known to produce optimal seeding.^52,53^ Mean length of sonicated fibril size varied between 35-43.6 nm for each batch of PFFs prepared (Figure 2).

**Figure 2:**
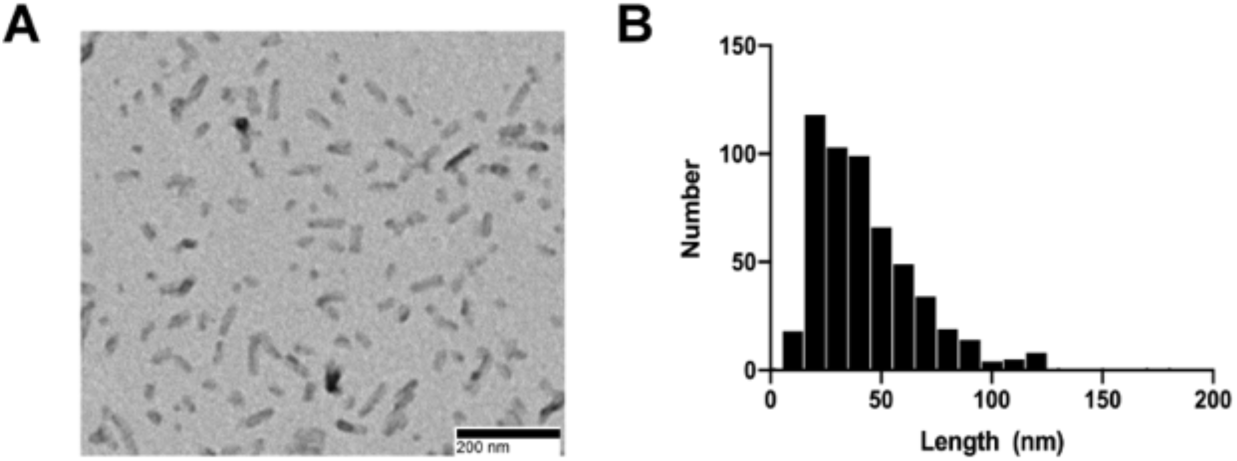
Sonicated α-syn PFFs. A) Representative TEM image of sonicated α-syn PFFs. B) Frequency distribution of fibril length.

### Intrastriatal injections of α-syn PFFs

Surgeries were performed as previously described, with slight modifications.^38^ Prior to surgery, mice were anesthetized with isoflurane. After anesthesia, 2.5 μl of liquid was unilaterally injected into the dorsal medial striatum using the following coordinates relative to bregma: anterior-posterior = 1.6 mm, medial-lateral = 2.0 mm, and dorsal ventral = −2.6mm. Injections were performed using pulled glass needles attached to 10 μl Hamilton syringes at a flow rate of 0.5 μl/minute. At the end of the injection, the needle was left in place for one minute, withdrawn 0.5 mm, left in place for an additional two minutes to avoid displacement of PFFs, and then completely retracted. Unilateral injections consisted of PBS (saline control) or 2 μg/μl α-syn PFFs (5 μg of total protein). During surgeries, PBS and PFFs were kept at room temperature. Post-surgery, animals received an analgesic (1 mg/kg of sustained release buprenorphine, subcutaneous administration) and were monitored closely until they recovered from anesthesia. In the three days following recovery, animals undergoing surgery were monitored daily for adverse outcomes.

### Motor behavior assessment (challenging beam)

The challenging beam was used to test motor performance and coordination. This test has been shown to be sensitive in detecting sensorimotor deficits in toxicant, α-syn, and genetic mouse models of PD with nigrostriatal dysfunction or neurodegeneration.^54–58^ Briefly, prior to beam training and testing, mice were acclimated to the behavior room for one hour. All behavioral experiments started at least one hour into the wake (dark) cycle of the mice. The plexiglass beam consisted of four 25 cm sections of gradually decreasing widths (3.5 cm, 2.5 cm, 2.0 cm, and 0.5 cm) and was assembled into a one-meter-long tapered beam. The home cage was placed at the end of the narrowest section to encourage mice to walk the length of the beam into their home cage. Mice were trained for two days on the tapered beam and received five trials each day. On the day of the test, the beam was made more challenging by placing a mesh grid (squares = 1 cm^2^) over the beam. The grid corresponded to the width of each beam section and created an ∼1 cm distance between the top of the grid and the beam. This allowed for the visualization of limb slips through the grid. On the day of the test, each mouse was videotaped for 5 trials. All mice were tested at baseline (prior to PFF injections) and at 4 and 6 months post-PFF injection. Videos were scored by trained raters blinded to experimental condition and with an inter-rater reliability of at least 90%. Raters scored the following outcome measures: time to traverse the beam, number of steps, and errors. An error was defined as a limb slip through the mesh grid during a forward movement.^54^ Each limb accounted for its own error (e.g. 2 slips in 1 forward movement = 2 errors). The mean of the 5 trials was used for analysis. For analysis, data were stratified by time point because experiments were not powered for longitudinal analysis.

### Motor behavior assessment (rotarod)

Rotarod testing was performed as previously described.^38^ Prior to rotarod training and testing, mice were acclimated to the behavior room for one hour. All behavioral experiments started at least one hour into the wake (dark) cycle of the mice. For training, each mouse received 3 practice trials with at least a 10-minute interval between each trial. For each trial, mice (n=10 per group) were placed on a rotarod with speed set at 5 rpm for 60 seconds. If an animal fell off, it was placed immediately back on the rotarod. Testing occurred 24 hours after training. For testing, the rotarod apparatus was set to accelerate from 4 to 40 rpm over 300 seconds and the acceleration was initiated immediately after an animal was placed on the rotarod. Each mouse underwent 3 trials with an inter-trial interval of at least 15 minutes. Males were always tested before the females, and the rods were cleaned between the trials to prevent cross-scents interfering with performance. All mice were tested at baseline (before receiving PFF injections) and at 4 and 6 months after PFF injections. The mean latency to fall in all 3 trials was used for analysis. For analysis, data were stratified by time point because experiments were not powered for longitudinal analysis.

### Tissue collection

All animals were euthanized by pentobarbital overdose and intracardially perfused with 0.9% phosphate-buffered saline. At the 1-month time point, saline perfusion was followed by perfusion with cold 4% paraformaldehyde (PFA) in phosphate-buffered saline and whole brains were extracted and post-fixed in 4% PFA for 24 hours and placed into 30% sucrose in phosphate-buffered saline for immunohistochemistry (IHC) at 4°C. For 2- and 6-month time points, brains were extracted after phosphate-buffered saline perfusion and rostral portions of each brain were flash frozen in 2-methylbutane on dry ice and stored at −80°C until use for HPLC. The caudal portions of each brain were post-fixed in 4% PFA for 24 hours at 4°C and placed into 30% sucrose for IHC.

### HPLC

Striatal tissue punches (1mm x 2mm) were collected from the dorsal striatum on a cryostat and sonicated in 200 µl of an antioxidant solution (0.4 N perchlorate, 1.34 mm EDTA, and 0.53 mm sodium metabisulfite). A 10 µl aliquot of the sonicated homogenate was removed into 2% SDS for BCA protein assay (Pierce). Remaining samples were clarified by centrifugation at 10,000 rpm for 10 minutes. Deproteinized supernatants were analyzed for levels of DA, HVA and DOPAC using HPLC. Samples were separated on a Microsorb MV C18 100–5 column (Agilent Technologies) and detected using a CoulArray 5200 12-channel coulometric array detector (ESA) attached to a Waters 2695 Solvent Delivery System (Waters) using the following parameters: flow rate of 1 ml/min; detection potentials of 25, 85, 120, 180, 220, 340, 420 and 480 mV; and scrubbing potential of 750 mV. The mobile phase consisted of 100 mm citric acid, 75 mM Na_2_HPO_4_, and 80 μm heptanesulfonate monohydrate, pH 4.25, in 11% methanol. Sample values were calculated based on a six-point standard curve of the analytes. Data were quantified as ng/mg protein.

### Western blotting

Western blots for α-syn, DAT and VMAT2 were carried out as previously described.^33,39,42,59–61^ Striatal tissue punches were homogenized in tissue lysis buffer (Tris buffered saline with 1% SDS) and centrifuged at 1150 x g for 5 minutes at 4°C. Protein levels were quantified by BCA protein assay. Samples were diluted with appropriate homogenization buffer, NuPage LDS sample buffer, and 100 mM DTT. For each sample, 10 µg of total protein for α-syn and 20 µg of total protein for DAT and VMAT2 was loaded onto a NuPage 10% Bis-Tris gel. Samples were co-blotted with dilution standards. Samples were subjected to PAGE and electrophoretically transferred to polyvinylidene difluoride membranes (ThermoFisher 88520). For α-syn detection only, immediately following the transfer, proteins were fixed to the membrane by incubating the membrane in 0.4% paraformaldehyde for 30 minutes at room temperature. After fixation, the membrane was incubated in REVERT staining solution (Li-Cor Biosciences, 926-11021) for 5 minutes and imaged on a Li-Cor Odyssey CLx for quantification of total protein. Non-specific sites were blocked with Odyssey Blocking Buffer (LI-COR Biosciences, 927-50003), and membranes were then incubated overnight in the appropriate primary antibody: α-syn (BD 610787, 1:500), DAT (Millipore Sigma MAB369, 1:1000) or VMAT2 (Miller Lab, 1:10,000).^61^ Primary antibody was prepared in 0.01% Tween in Odyssey Blocking Buffer. Primary antibody binding was detected with the appropriate secondary antibody (α-syn: IRDye 800CW Goat anti-Mouse, Li-Cor 926-32210; DAT: IRDye 800CW Goat anti-Rat, Li-Cor 926-32219, 1:10,000; VMAT2: IRDye 800CW Goat anti-Rabbit, Li-Core 926-32219) and imaged on a Li-Cor Odyssey CLx. Quantifications of both total protein and bands of interest were performed in Image Studio Lite Version 5.2. Intensities were calibrated to co-blotted dilutional standards and normalized to total protein.

### Taqman Array Cards

RNA isolation was performed using the RNeasy Lipid Tissue Mini Kit (Qiagen), with minor modifications to improve RNA yield. First, tissue was homogenized in 200 µl cold Qiazol lysis reagent. Second, after homogenization, an additional 800 µl of Qiazol lysis reagent was added to each tissue sample followed by 200 µl of chloroform. Third, to facilitate separation of the RNA containing aqueous layer, samples were centrifuged in Phasemaker tubes. Finally, the optional DNase digestion step was included to improve purity of isolated RNA. RNA was eluted in 50 µl RNase-free water, and RNA yield and purity were both assessed using the Agilent RNA 6000 Pico Reagents with the Agilent 2100 Bioanalyzer System (Agilent Technologies). RIN scores were between 7.7 and 8.6 for all samples. Isolated RNA was stored at −80°C.

cDNA synthesis was performed according to directions supplied with Superscript IV VILO master mix (Life Technologies). 750 ng of RNA input was used per reaction. Reactions also included 1 µl of RNaseOUT (ThermoFisher). cDNA was stored at −20°C until use. qPCR reactions were prepared according to the manufacturer’s protocol using Taqman Fast Advanced Mastermix (ThermoFisher). The TaqMan Array Mouse Immune Panel (ThermoFisher) was run on a Viia7 Real-Time PCR instrument (ThermoFisher) according to kit instructions.

The 2^-ΔΔCt^ method was performed on the Taqman array data to estimate relative changes in gene expression by dieldrin exposure.^62^ ΔCt was calculated as follows: ΔCt = Ct (mean of target gene) – Ct (mean of housekeeping genes). Four housekeeping genes were used to calculate the housekeeping gene mean: *Actb, Gusb, Hprt1*, and *Gapdh*. We excluded *18S rRNA* as a housekeeping gene because the assay failed in some samples. The ΔΔCt value for each gene was calculated as follows: ΔCt (dieldrin group) - ΔCt (control group). Fold change was calculated as follows: fold change = 2^−ΔΔCt.^. To test for differential gene expression between dieldrin and control brains, we ran Welch’s two-sample t-tests comparing ΔCt values for each gene in the two experimental groups. Significance level for t-tests was set at p< 0.05. All gene expression analyses were stratified by sex. Lists of significant differentially expressed genes in male and female mouse brains were input into the STRING network tool to test for known protein-protein interactions. In STRING, the meaning of network edges was set to represent molecular action; otherwise, we used default settings.

### Immunohistochemistry

Fixed brains were frozen on a sliding microtome and sliced at 40 μm. Free-floating sections were stored in cryoprotectant (30% sucrose, 30% ethylene glycol, 0.05M PBS) at −20°C. A 1:4 series was used for staining. Nonspecific staining was blocked with 10% normal goat serum. Sections were then incubated overnight in appropriate primary antibody: p-syn (Abcam, Ab184674, 1:10,000) or TH (Millipore, AB152, 1:4,000). Primary antibodies were prepared in Tris-buffered saline with 1% NGS/0.25% Triton X-100. Sections were incubated with appropriate biotinylated secondary antibodies at 1:500 (anti-mouse, Millipore AP124B or anti-Millipore AP132B), followed by Vector ABC standard detection kit (Vector Laboratories PK-6100). Visualization was performed using 0.5 mg/ml 3,3’ diaminobenzidine (DAB, Sigma-Aldrich) for 30-60 seconds at room temperature and enhanced with nickel. Slides stained for p-syn were counter-stained with Cresyl violet. Slides were dehydrated before cover-slipping with Cytoseal (Richard-Allan Scientific) and imaged on a Nikon Eclipse 90i microscope with a QICAM camera (QImaging) and Nikon Elements AR (version 4.50.00).

### Quantification of p-syn inclusion-bearing neurons in the SNpc

Total enumeration of neurons containing p-syn was performed as previously described using StereoInvestigator (MBF Bioscience).^50^ During counting, the investigator was blinded to the treatment groups. Sections containing the SNpc (1:4 series) were used for all counts. Contours were drawn around the SNpc using the 4x objective. A 20x objective was used to identify the stained inclusions. All neurons containing p-syn within the contour were counted and total counts were multiplied by four to estimate the total number of neurons in each animal with inclusions. Samples injected with PFFs that did not have any p-syn pathology were excluded as missed injections.

### Stereology

TH and NeuN neuron counts were estimated by unbiased stereology with StereoInvestigator (MBF Bioscience) using the optical fractionator probe, as described previously.^63–65^ Briefly, sections containing the SNpc (1:4 series) were used for all counts. In all cases, the investigator was blinded to the treatment groups. Contours around the SNpc were drawn using the 4x objective and counting was done using a 60X oil immersion objective. The following settings were used for TH counting: grid size = 195 μm x 85 μm, counting frame = 50 μm x 50 μm, guard zone = 3 μm, and optical dissector height = 23 μm. The following settings were used for NeuN counting: grid size = 260 μm x 320 μm, counting frame = 50 µm x 50 µm, guard zone = 2.5 μm, and optical dissector height = 25 μm. Section thickness was measured every third counting frame, with an average measured thickness of 29 μm. Labeled neurons within the counting frame were counted while optically dissecting the entire section through the z-axis. Variability was assessed with the Gundersen coefficient of error (≤0.1).

### Data analysis and statistics

Statistical analysis of all data and graphing were performed using either GraphPad Prism or R (version 3.5.3). All analyses were stratified by sex. All two-group comparisons (total enumeration of p-syn, baseline behavior, Taqman array cards and western blots) were performed using an unpaired Welch’s t-test. Stereology, HPLC results and post-PFF behavior were compared by two-way ANOVA followed by Sidak’s multiple comparisons tests. Linear regression was performed to test for associations between HPLC results and behavioral outcomes using the lm() function in R. Pregnant dams were the experimental unit for all analyses and all pups for each outcome came from independent litters. All data are shown as mean +/-95% CI. Results of two-way ANOVA are included in figure legends and significant results of Sidak post-tests are indicated on graphs.

## Results

### Dieldrin exposure exacerbates PFF-induced deficits in motor behavior

Animals were tested on challenging beam and rotarod prior to PFF injections (baseline), and at 4 months and 6 months post-PFF injections. On the rotarod, we observed no PFF- or dieldrin-induced deficits in the latency to fall at any time point (Supplementary Figure 1). On challenging beam, a sensorimotor test that assesses fine motor coordination and balance in mice, the biggest differences were detected at the 6 month time point (Figure 3).^54–58^ Results from each timepoint are described below.

**Figure 3:**
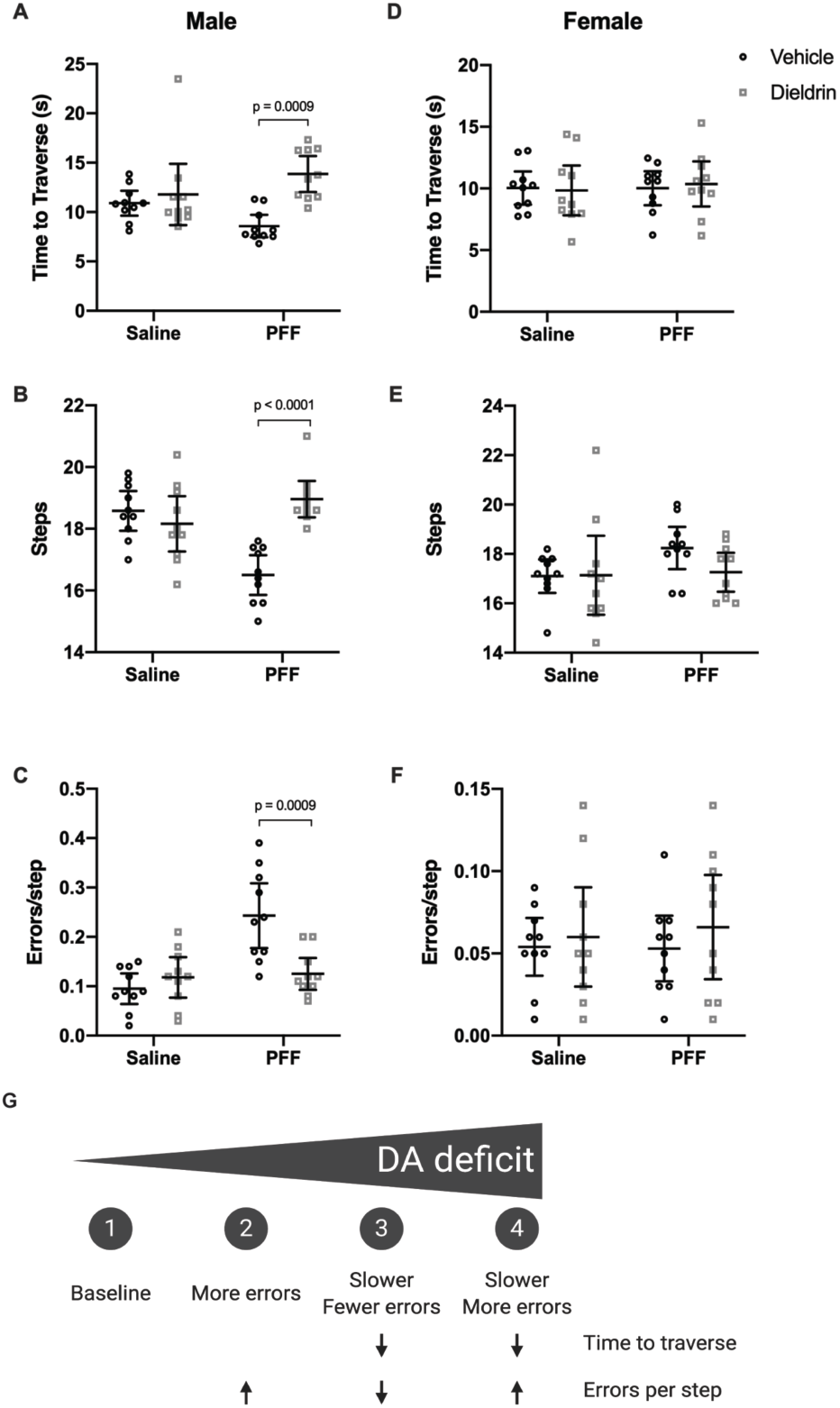
Dieldrin exacerbates PFF-induced motor deficits on challenging beam in male animals only. Six months after PFF- injection, motor behavior was assessed on challenging beam in male (A-C) and female (D-F) animals (n = 10 per group). Time to traverse (A,D), steps across the beam (B,E) and errors per step (C,F) were scored. A) Time to traverse at 6 months after PFF injection in male animals (two-way ANOVA: PFF, p = 0.8914; dieldrin, p = 0.0013; interaction, p = 0.0171). Sidak post-tests showed a significant dieldrin-related increase in time to traverse in PFF injected animals (vehicle:PFF vs dieldrin:PFF animals, p = 0.0009). B) Steps at 6 months after PFF injection in male animals (two-way ANOVA: PFF, p = 0.0023; dieldrin, p = 0.0469; interaction, p < 0.0001). Sidak post-tests showed a significant dieldrin-related increase in steps in PFF-injected animals (vehicle:PFF vs dieldrin:PFF animals, p < 0.0001), as well as a significant effect of PFF in vehicle exposed animals (vehicle:saline vs vehicle:PFF, p = 0.0002). C) Errors per step at 6 months post-PFF injection (two-way ANOVA: PFF, p = 0.0004; dieldrin, p = 0.0215; interaction, p = 0.0010). Sidak post-tests showed a significant dieldrin-related decrease in errors per step in PFF-injected animals (vehicle:PFF animals vs dieldrin:PFF animals, p=0.0009), as well as a significant effect of PFF in vehicle exposed animals (vehicle:saline vs vehicle:PFF, p < 0.001). D-F) In female animals, all results were non-significant. G) Schematic illustrating the progression of motor deficits as DA deficits become more severe. All data shown as mean +/- 95% CI with significant results of Sidak post-tests for dieldrin to vehicle comparisons indicated on graphs. All significant post-test results are reported in this legend.

**Supplementary Figure 1:**
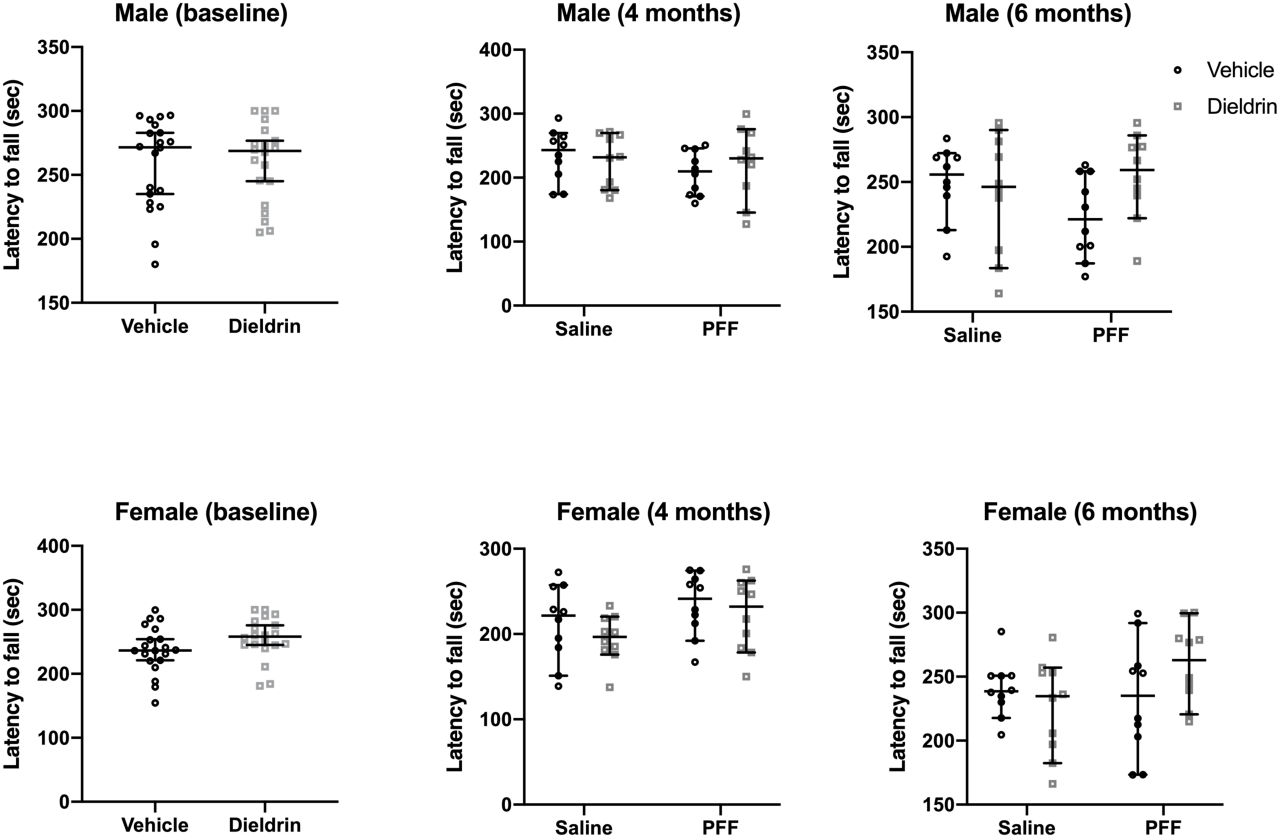
No PFF- or dieldrin-induced effects were observed on rotarod in male or female mice. Male and female mice (n = 20 per group at baseline, n = 10 per group at later time points) were tested on rotarod prior to PFF injections (Baseline) and at 4 and 6 months post-PFF injections. Latency to fall in both sexes at all time points, except females at 4 months, was unaffected by either PFF injection or dieldrin exposure. In females at 4 months, there was a significant PFF effect by two-way ANOVA (p = 0.0478), but no single comparison was significant by Sidak post-test. Data is shown as mean (95% CI)

#### Baseline

There were no significant differences in any of the outcome measures for either sex at baseline testing (Table 1).

**Table 1:** Challenging beam performance in male and female mice at baseline. Male and female mice (n = 20 per group) were tested on challenging beam prior to PFF injections. Data shown as mean (95% CI). Results of Welch’s t-tests to compare vehicle and dieldrin were all not significant.

| **Male** | **Vehicle** | **Dieldrin** |
| --- | --- | --- |
| *Time to traverse* | 11.60 (10.45-12.74) | 11.05 (9.892-12.21) |
| *Steps* | 20.01 (18.89-21.13) | 19.24 (18.82-19.66) |
| *Errors/Step* | 0.047 (0.036-0.058) | 0.050 (0.036-0.063) |
| **Female** |  |  |
| *Time to traverse* | 12.84 (11.52-14.16) | 12.85 (11.45-14.25) |
| *Steps* | 17.80 (17.43-18.17) | 16.47 (15.37-17.57) |
| *Errors/Step* | 0.0885 (0.07-0.10) | 0.095 (0.073-0.117) |

#### 4 months post PFF injection

In male animals, we observed no differences between the groups in time to traverse the beam (Table 2). However, dieldrin exposed animals (dieldrin:saline) made fewer steps across the beam than vehicle controls (vehicle:saline). PFFs had no effect on steps in vehicle exposed animals but caused an increase in steps in dieldrin exposed animals. In addition, there was a significant effect of PFF on errors per step at this time point. Post-tests did not identify a significant effect of dieldrin despite a higher error rate in dieldrin exposed animals (dieldrin:saline), but did identify a PFF-induced reduction in errors in dieldrin exposed animals.

**Table 2:**
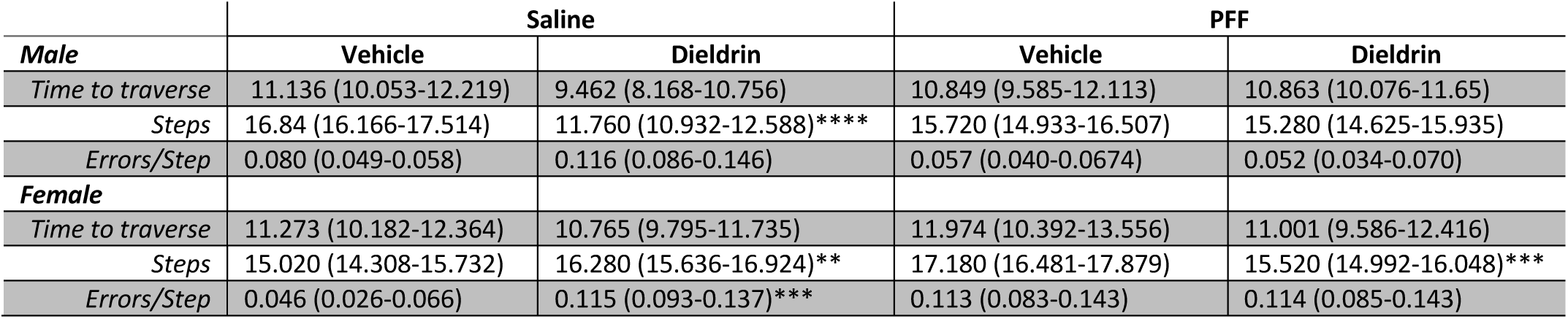
Challenging beam performance in male and female mice at 4 months post-PFF injection. Male and female mice (n = 10 per group) were tested on challenging beam 4 months after PFF injections. Two-way ANOVA with Sidak’s multiple comparison tests were performed. Results of Sidak post-tests between dieldrin exposed animals and corresponding vehicle controls are indicated on the table. All significant post-test results are included in this legend. *Males:* There were no significant differences in males on time to traverse. On steps, PFF, dieldrin and the interaction were all statistically significant by two-way ANOVA (PFF: p = 0.0008; dieldrin, p < 0.0001; Interaction: p < 0.0001). Sidak post-tests shows a significant effect of dieldrin in saline-injected animals (vehicle:saline vs dieldrin:saline, p < 0.0001) but not in PFF-injected animals, as well as an effect of PFF in dieldrin exposed animals (dieldrin:saline vs dieldrin:PFF, p <0.0001), but no effect of PFF in vehicle animals. On errors/step, there was a significant effect of PFF by two-way ANOVA (PFF: p = 0.003; dieldrin, p = 0.1657; interaction, p = 0.0695). Sidak post-tests showed no significant effect of dieldrin, but did identify a PFF effect in dieldrin exposed animals (dieldrin:saline vs dieldrin:PFF, p = 0.0012). *Females:* There were no significant differences in time to traverse. On steps, PFF and the interaction were statistically significant (PFF: p = 0.0199; dieldrin, p = −.4907; interaction: p < 0.0001). Sidak post-tests revealed a significant effect on dieldrin in both saline- and PFF-injected animals (vehicle:saline vs dieldrin:saline, p = 0.0222; vehicle:PFF vs dieldrin: PFF, p = 0.0014), as well as a PFF-effect in vehicle exposed animals but not dieldrin exposed animals (vehicle:saline vs vehicle:PFF, p =< 0.0001; dieldrin:saline vs dieldrin:PFF, p =0.3510). On errors/step, PFF, dieldrin and the interaction were all statistically significant by two-way ANOVA (PFF: p = 0.0061; dieldrin, p = 0.0038; Interaction: p = 0.0048). Sidak post-tests revealed a significant effect of dieldrin in saline-injected animals (vehicle:saline vs dieldrin:saline, p = 0.0007), as well as a PFF-related increase in errors in vehicle animals (vehicle:saline vs vehicle:PFF, p = 0.0011).

In female animals, there were also no differences between the groups in time to traverse (Table 2). Dieldrin exposed animals (dieldrin:saline) made more steps across the beam than their vehicle control, while in PFF-injected animals, dieldrin exposed animals (dieldrin:PFF) made fewer steps than their vehicle control (vehicle:PFF). PFFs also induced a significant increase in steps in vehicle exposed animals. Control animals (vehicle:saline) made fewer errors per step than the other groups at this time point and fewer errors than they did at baseline; the other treatment groups made similar errors per step than they did at baseline (Table 2,Supplementary Figure 2F).

#### 6 months post PFF injection

In male animals, dieldrin exposure was associated with a 40% increase in time to traverse in PFF-injected animals, but we observed no effect of PFF alone (Figure 3A). For steps, PFF-injection alone caused a significant decrease in steps, and there was a significant dieldrin-associated increase in PFF-injected animals, with dieldrin exposed animals showing a 13% increase in steps (Figure 3B). For errors per step, there was a robust PFF-related increase in the vehicle exposed animals that was not observed in dieldrin exposed animals that received PFF injections (Figure 3C). While these dieldrin:PFF animals did not make as many errors as the vehicle:PFF group, they did make more errors compared to their baseline performance (Supplementary Figure 2C). In contrast, at this timepoint in females, there were no differences between the groups in any of the outcomes (Figure 3D-F).

**Supplementary Figure 2:**
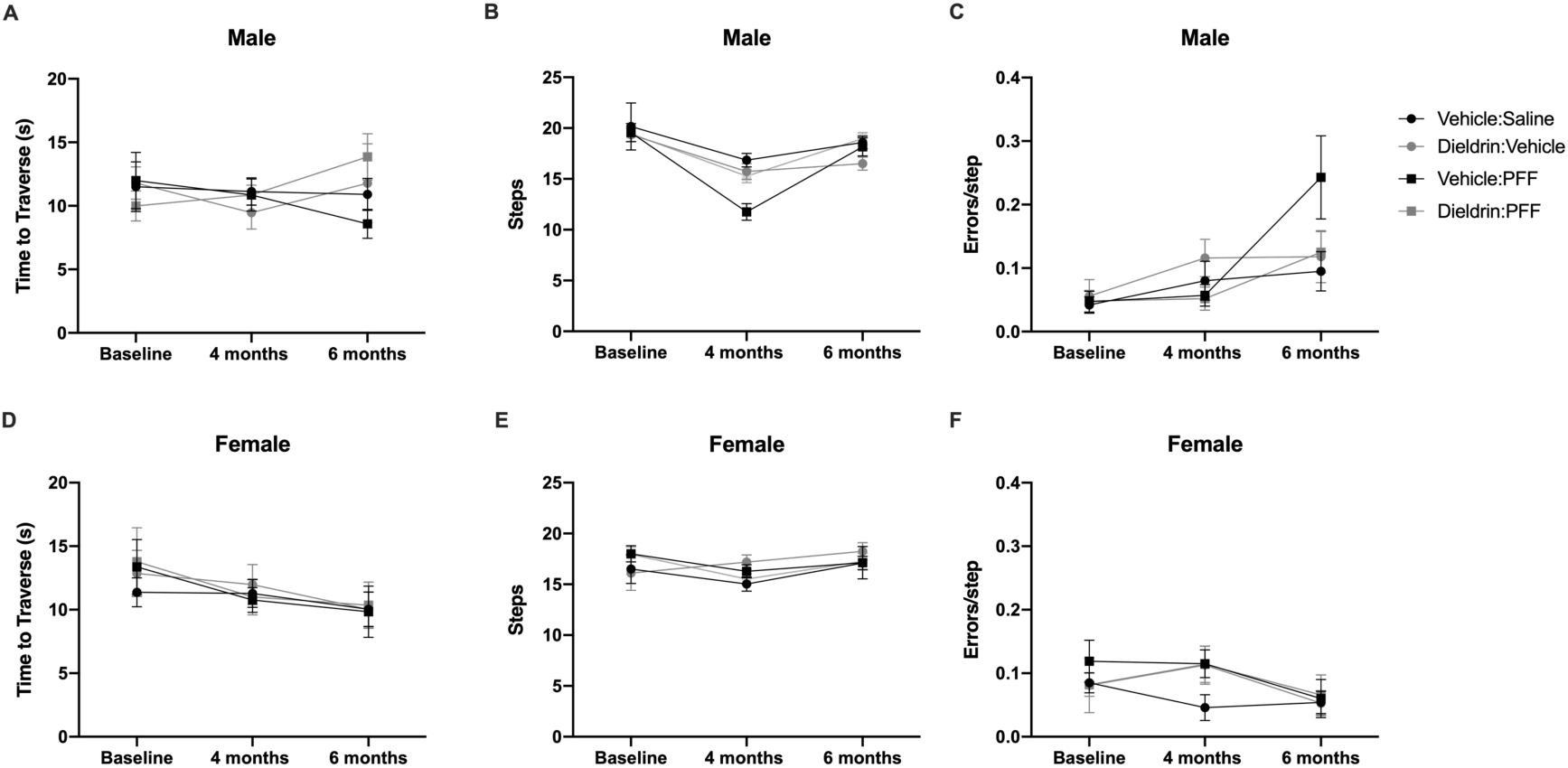
Behavioral data at all time points. All challenging beam data presented longitudinally to highlight changes over time. A-C) Male animals; D-F) Female animals; A,D) Time to traverse; B,D) Steps; C,F) Errors per step. Statistical analysis was stratified by time point since we lacked statistical power for a longitudinal analysis.

### Dieldrin exposure does not increase PFF-induced pSyn aggregation

In mice, phosphorylated α-syn (p-syn) aggregates accumulate progressively unti 2 months after PFF injections in the substantia nigra (SN) pars compacta, evolving from pale cytoplasmic inclusions 1 month post-PFF injection to dense perinuclear Lewy body-like inclusions by 3 and 6 months post-PFF injection.^38^ To determine whether developmental dieldrin exposure increases the propensity for α-syn to aggregate, we quantified the number of p-syn-containing neurons in the ipsilateral SN at 1 and 2 months post-PFF injection in mice developmentally exposed to dieldrin or vehicle. Our results showed that developmental dieldrin exposure had no effect on the number of p-syn-containing neurons in the ipsilateral nigra at 1 or 2 months post-PFF injection in male or female animals (Figure 4). The number of psyn-containing neurons was similar between males and females. Consistent with previous observations, we observed no neurons containing p-syn inclusions in the contralateral SN.

**Figure 4:**
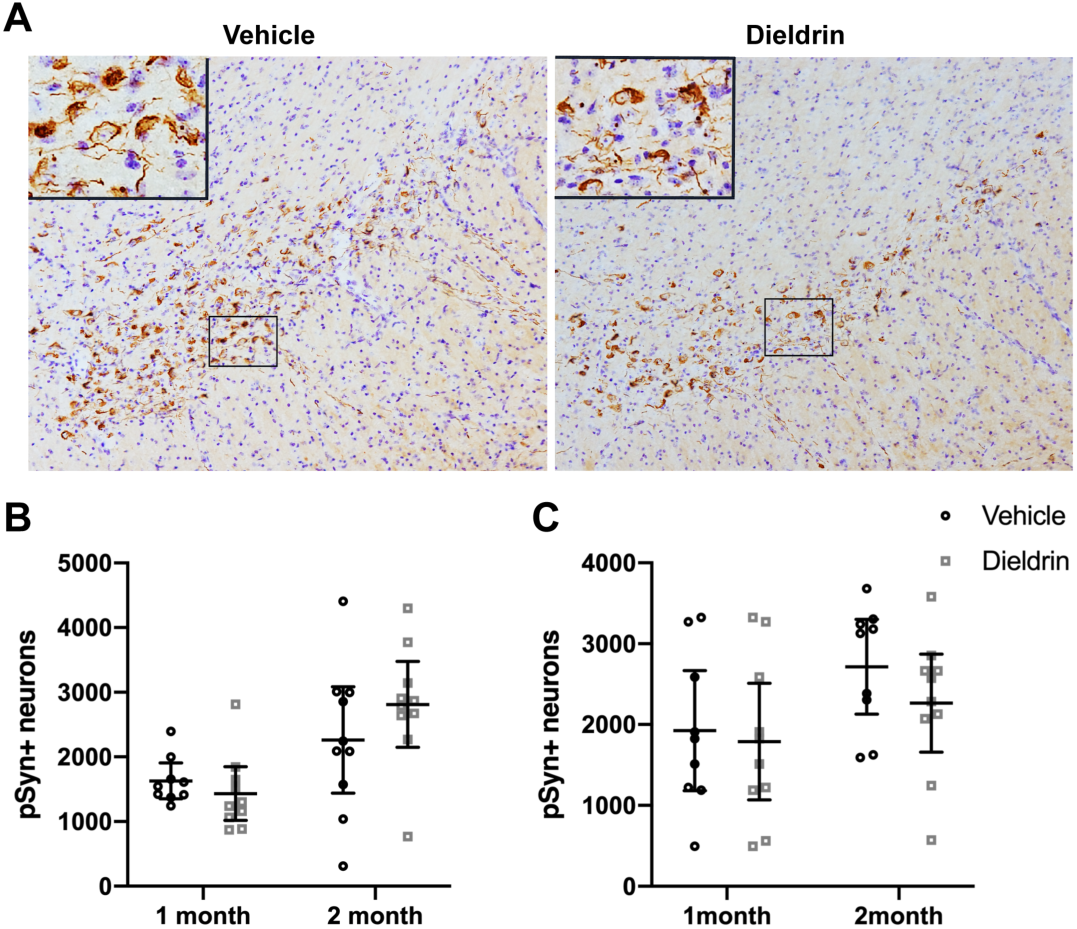
Developmental dieldrin exposure does not affect the propensity of p-syn to accumulate in the SN. A) Representative images of psyn immunohistochemistry from the identical coronal levels through the ipsilateral SN in male animals 2 months after intrastriatal PFF injection. B) Total enumeration of psyn-containing neurons in ipsilateral SN in male animals (n = 9 for 1 month vehicle due to seeding failure in 1 animal; n = 10 in all other groups; unpaired t-test with Welch’s correction: 1 month, p = 0.1919; 2 month, p = 0.1272). C) Total enumeration of psyn-containing neurons in ipsilateral SN in female animals (n = 9 for 1 month vehicle due to seeding failure in 1 animal; n = 10 in all other groups; unpaired t-test with Welch’s correction: 1 month, p = 0.4712; 2 month, p = 0.1195). Data shown as mean +/- 95% CI.

### Dieldrin exposure exacerbates PFF-induced increases in DA turnover

To test if dieldrin exacerbated PFF-induced decreases in striatal DA levels, we measured DA and two of its metabolites, DOPAC and HVA, in ipsilateral and contralateral dorsal striatum by HPLC at 2 and 6 months post-PFF injections. Consistent with previous results, we showed PFF-induced deficits in DA levels (∼45% loss at 6 months) in the ipsilateral dorsal striatum of male animals at both time points, but this loss was not exacerbated by prior dieldrin exposure (Figure 5A, Supplementary Figure 3A).^38^ We also showed, for the first time, that female mice exhibit a PFF-induced loss of striatal DA (∼40% loss at 6 months) at both time points (Figure 5F, Supplementary Figure 3F). In both male and female animals, we observed PFF-induced deficits in DOPAC and HVA at the two measured time points, but there was no exacerbation of this loss by dieldrin in either sex (Figure 5B,C,G,H, Supplementary Figure 3B,C,G,H). These PFF-induced deficits were also seen at 2 months post-PFF injection (Supplementary Figure 3A-C, F-H). Dieldrin had no effect on these outcome measures at 2 months in male animals. In female animals, dieldrin significantly reduced the PFF-induced loss of HVA levels, suggesting that the progression of striatal dysfunction is slower in female animals (Supplementary Figure 3H).

**Figure 5:**
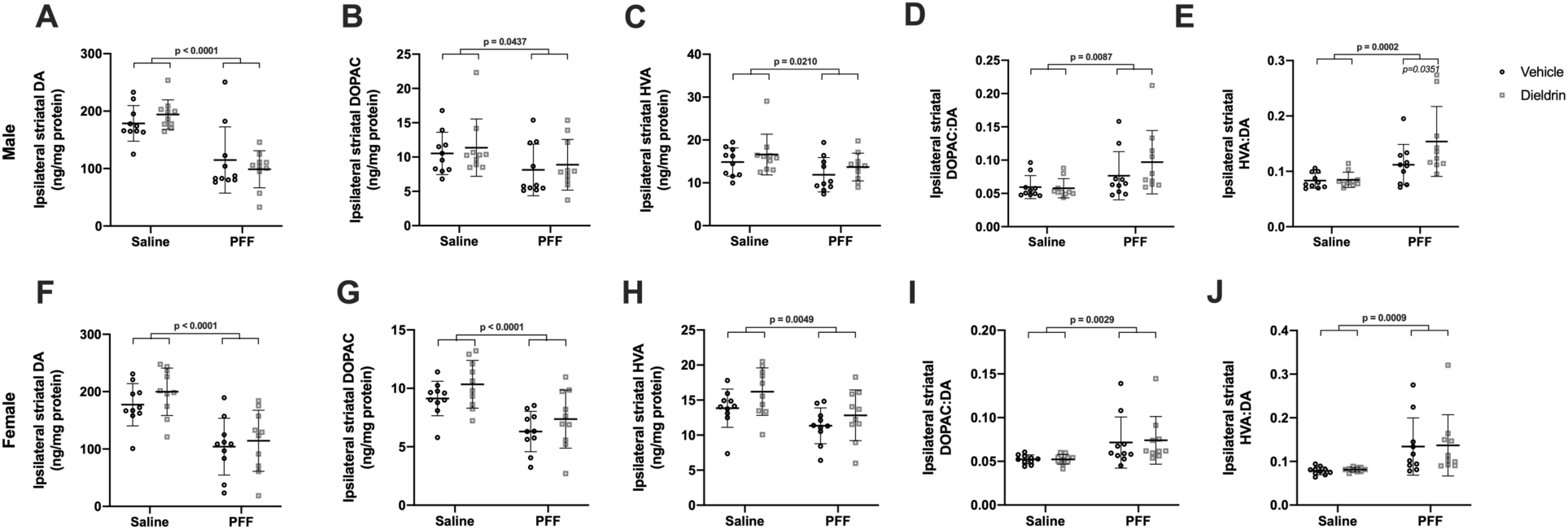
Developmental dieldrin exposure exacerbates PFF-induced increases in DA turnover in male animals only 6 months after PFF injection. Levels of DA, DOPAC, and HVA in the ipsilateral dorsal striatum were measured 6 months post-PFF injection by HPLC in male (A-E) and female (F-J) animals (n = 10 per group). A) PFF-induced loss of DA levels in ipsilateral dorsal striatum in male animals (two-way ANOVA: PFF, p < 0.0001; dieldrin, p = 0.9788, interaction, p = 0.2052). B) PFF-induced loss of DOPAC levels in ipsilateral dorsal striatum in male animals (two-way ANOVA: PFF, p = 0.0437; dieldrin, p = 0.9671; interaction, p = 0.5027). C) PFF-induced loss of HVA levels in ipsilateral dorsal striatum in male animals (two-way ANOVA: PFF, p = 0.0210; dieldrin, p = 0.1558; interaction, p = 0.9847). D) PFF-induced increase in DOPAC:DA ratio in ipsilateral dorsal stratum of male animals (two-way ANOVA: PFF, p = 0.0087; dieldrin, p = 0.3607; interaction, p = 0.2814). E) HVA:DA ratio in ipsilateral dorsal striatum of male animals (two-way ANOVA: PFF, p = 0.0002; dieldrin, p = 0.0786; interaction, p = 0.0967). Sidak post-tests showed a significant effect of dieldrin in PFF injected animals (vehicle:PFF vs.dieldrin:PFF, p = 0.0351), but not saline injected animals. F) PFF-induced loss of DA levels in ipsilateral dorsal striatum of female animals (two-way ANOVA: PFF, p < 0.0001; dieldrin, p = 0.2667; interaction, p = 0.6746). G) PFF-induced loss of DOPAC levels in ipsilateral dorsal striatum in female animals (two-way ANOVA: PFF, p < 0.0001; dieldrin, p = 0.0654; interaction, p = 0. 8994). H) PFF-induced loss of HVA levels in ipsilateral dorsal striatum of female animals (two-way ANOVA: PFF, p = 0.0049; dieldrin, p = 0.0565; interaction, p = 0.6614). I) PFF-induced increase in DOPAC:DA ratio in ipsilateral dorsal striatum of female animals (two-way ANOVA: PFF, p = 0.0029; dieldrin, p = 0.8397; interaction, p = 0.8502). J) PFF-induced increase in HVA:DA ratio in ipsilateral dorsal striatum of female animals (two-way ANOVA: PFF, p = 0.0009; dieldrin, p = 0.8577; interaction, p = 0.9994). Data shown as mean +/- 95% CI with significant results of two-way ANOVA indicated on graphs in bold and of Sidak post-tests for dieldrin to vehicle comparisons indicated in italics. Except where indicated (E), all measures had significant PFF-induced deficits, but no significant effect of dieldrin.

To investigate the effects of dieldrin exposure and PFFs on DA turnover, we calculated ratios of DOPAC and HVA to DA. At both time points, in both sexes, we observed PFF-induced increases in both the DOPAC:DA and HVA:DA ratios, indicative of increased DA turnover and deficits in DA packaging. In male animals at 6 months post-PFF injection only, this increase in HVA:DA ratio was further exacerbated by prior dieldrin exposure (Figure 5E). This dieldrin-induced exacerbation was not observed in females animals at 6 months post-PFF injection or at 2 months post-PFF injection in either sex (Figure 5J, Supplementary Figure 3E,J).

As expected, levels of DA and its metabolites remained unchanged following PFF injection in the contralateral striatum and dieldrin exposure alone had no effect on DA levels, DA metabolites, or DA turnover in the contralateral striatum (Supplementary Figure 4).

**Supplementary Figure 3:**
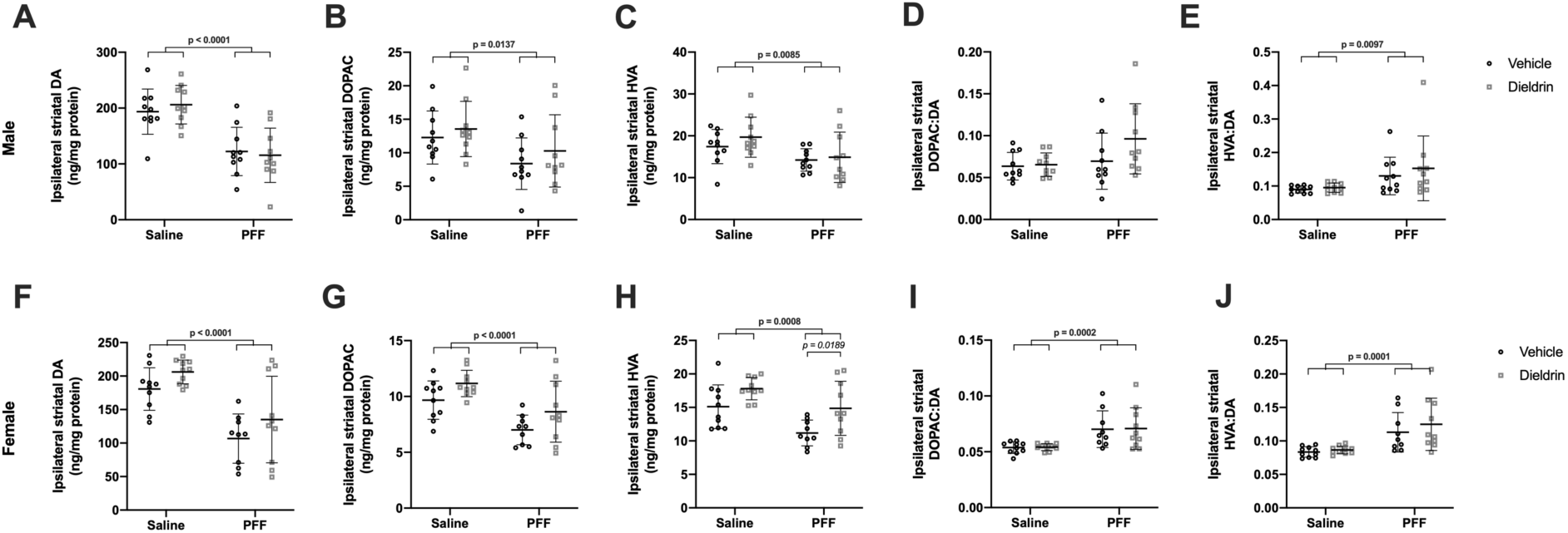
HPLC results at 2 months post-PFF injections. Levels of DA, DOPAC and HVA in ipsilateral dorsal striatum were measured 2 months post-PFF injection by HPLC in male (A-E) and female (F-J) animals (n = 10 per group; n = 9 for vehicle:PFF female group, one animal was excluded due to PFF seeding failure based on psyn counts). A) PFF-induced loss of DA levels (two-way ANOVA: PFF, p < 0.0001; dieldrin, p = 0.8310; interaction, p = 0.4696). B) PFF-induced loss of DOPAC levels (two-way ANOVA: PFF, p = 0.0137; dieldrin, p = 0.2607; interaction, p = 0.8189). C) PFF-induced loss of HVA levels (two-way ANOVA: PFF, p = 0.0085; dieldrin, p = 0.3247; interaction, p = 0.5916). D) No significant effect of PFF or dieldrin on DOPAC:DA ratio (two-way ANOVA: PFF, p = 0.0810; dieldrin, p = 0.1144; interaction, p=0.1597). E) PFF-induced increase in HVA:DA ratio (two-way ANOVA: PFF, p = 0.0097; dieldrin, p = 0.4414; interaction, p = 0.6346). F) PFF-induced loss of DA levels (two-way ANOVA: PFF, p < 0.0001; dieldrin, p = 0.0508; interaction, p = 0.9145). G) PFF and dieldrin effects on DOPAC levels (two-way ANOVA: PFF, p < 0.0001; dieldrin, p = 0.0129; interaction, p = 0.9125). H) PFF and dieldrin effects on HVA levels (two-way ANOVA: PFF, p = 0.0008; dieldrin, p = 0.0017; interaction, p = 0.6033). Sidak post-tests show a dieldrin-induced increase in HVA in PFF-injected animals (vehicle:PFF vs dieldrin:PFF: p = 0.0189). I) PFF-induced increase in DOPAC:DA ratio (two-way ANOVA: PFF, p = 0.0002; dieldrin, p = 0.9054; interaction, p=0.9842). J) PFF-induced increase in HVA:DA ratio (two-way ANOVA: PFF, p = 0.0001; dieldrin, p = 0.3585; interaction, p = 0.5825). Data shown as mean +/- 95% CI with significant results of two-way ANOVA indicated on graphs in bold and of Sidak post-tests indicated in italics. Except where indicated (H), Sidak post-hoc tests showed no significant effect of dieldrin exposure.

**Supplementary Figure 4:**
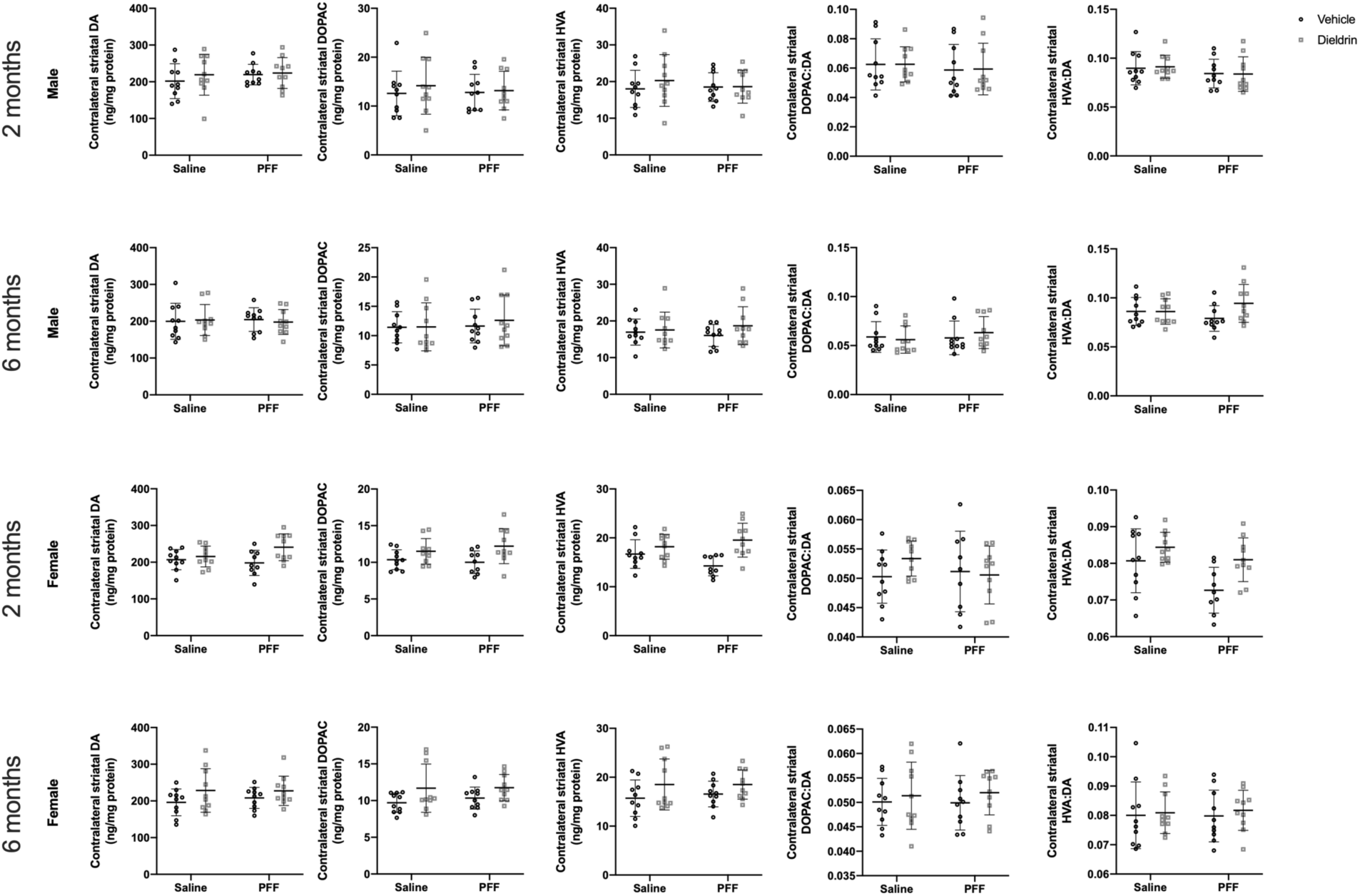
Contralateral HPLC at 2 and 6 months after PFF injections in male and female animals. As expected, there was no significant effect of PFF injection or dieldrin exposure on DA, DOPAC or HVA levels in male or female animals (n = 10 per group), at either timepoint. Data shown as mean +/- 95% CI.

The combination of male-specific deficits in motor behavior and DA turnover at 6 months post-PFF injection suggests that this dieldrin-induced exacerbation is due to increased synaptic deficits in the striatum. To further explore this potential link between DA turnover and the observed behavioral phenotype, we carried out linear regression for all male PFF-injected animals, regardless of dieldrin status, to determine if there is an association between DA turnover and behavioral outcome measures. We found a statistically significant association between HVA:DA ratio and time to traverse: the more severe the deficit in DA turnover, the more severe the behavioral deficit on time to traverse (Figure 6). Although we also observed a negative relationship between HVA:DA ratio and errors per step, this association was not statistically significant.

**Figure 6:**
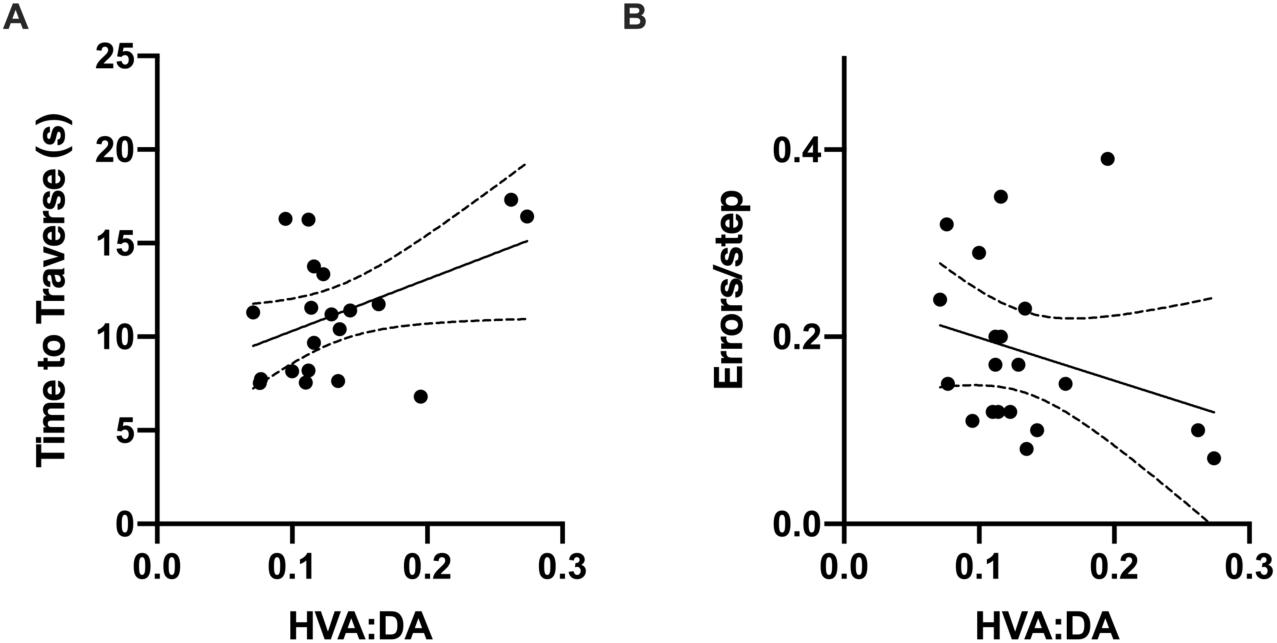
DA turnover is associated with behavioral phenotype. Linear regression was performed for all male PFF-injected animals with no additional covariates or interaction terms to explore associations between DA turnover and motor behavior outcomes. A) There was a significant positive association between DA turnover (HVA:DA) and time to traverse (beta coefficient = 27.651, p=0.0498). B) In contrast, the negative association between DA turnover (HVA:DA) and errors per step was not significant (beta coefficient = −0.46011, p-value = 0.248).

### Dieldrin exposure does not exacerbate PFF-induced loss of TH phenotype or neuronal loss in the substantia nigra

To determine if developmental dieldrin exposure exacerbates the PFF-induced loss of TH phenotype, we performed IHC for TH and estimated the number of TH^+^ neurons in the SN 6 months post-PFF injection by stereology. Consistent with prior results, we observed a ∼35% loss of TH^+^ neurons ipsilateral to the injection site in the SN 6 months after PFF injections (Figure 7A,D).^38^ Developmental dieldrin exposure did not significantly affect PFF-induced loss of TH^+^ neurons in male animals (Figure 7A,D).

**Figure 7:**
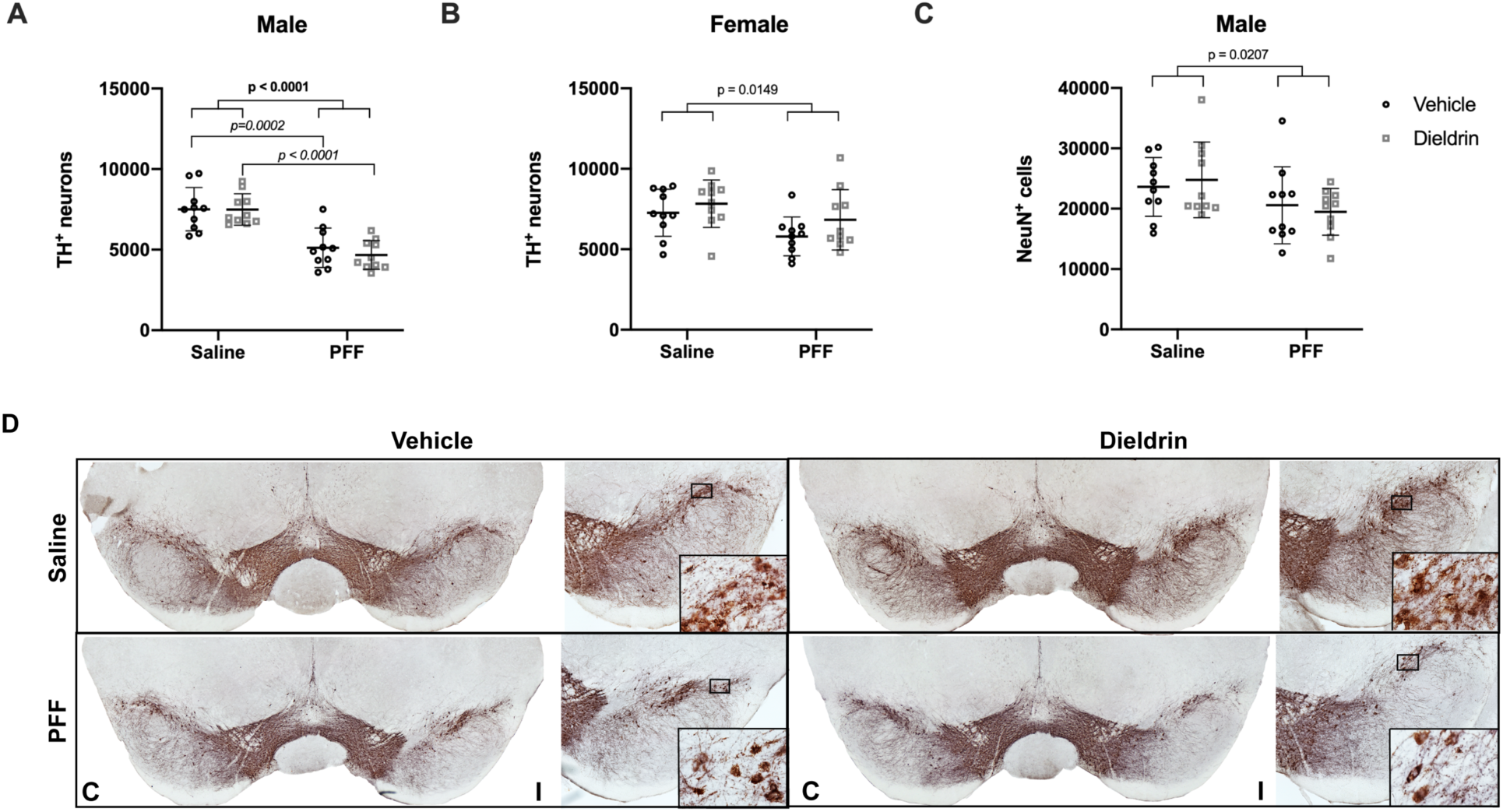
Dieldrin does not exacerbate the male-specific PFF-induced loss of ipsilateral nigral TH immunoreactive neurons. Number of TH^+^ neurons in the ipsilateral nigra was estimated by unbiased stereology. A) Ipsilateral nigral TH neuron counts in male animals (n = 10 per group) show a PFF-induced loss of TH^+^ neurons (two-way ANOVA: PFF, p < 0.0001; dieldrin, p = 0.5215; interaction, p = 0.5444). Sidak post-tests show no significant effect of dieldrin, but a significant effect of PFFs in vehicle and dieldrin exposed animals (vehicle:saline vs vehicle:PFF, p = 0.0002; dieldrin:saline vs dieldrin:PFF, p < 0.0001). B) Quantification of ipsilateral nigral NeuN counts in male animals (n = 10 per group) show a PFF-induced loss of NeuN (two-way ANOVA: PFF, p = 0.0207; dieldrin, p = 0.9823; interaction, p = 0.5133). Sidak post-tests show no significant effect in any individual comparison. C) Quantification of ipsilateral nigral TH counts in female animals (n = 10 per group) show a PFF effect (two-way ANOVA: PFF = 0.0149; dieldrin = 0.1061; interaction p = 0.6275). Sidak post-tests show no significant effect of PFFs or dieldrin. The only significant post-test was between dieldrin:saline and vehicle:PFF (p = 0.0304). D) Representative images from male animals of nigral TH immunohistochemistry. “C” and “I” indicate contralateral and ipsilateral sides. Data shown as mean +/- 95% CI with significant results of two-way ANOVA indicated on graphs in bold and of Sidak post-tests for dieldrin to vehicle comparisons indicated on graphs in italics. All significant post-test results are reported in this legend.

In contrast, in female animals, there was a significant effect of PFF on number of TH^+^ neurons, with a less than 20% loss of TH^+^ neurons ipsilateral to the injection site in the SN 6 months after PFF injections (Figure 7C). However, post-tests revealed no significant effect of dieldrin or PFF alone. As expected, there was no loss of TH^+^ neurons in the contralateral uninjected SN in either male or female animals (Supplementary Figure 5).

To assess whether the loss of TH immunoreactivity in PFF-injected male animals was accompanied by degeneration of these neurons, we performed IHC for NeuN in male animals and estimated the number of NeuN^+^ neurons in the SN by stereology. We observed a PFF-induced loss of NeuN^+^ neurons (∼20%) in the ipsilateral SN, with no effect of dieldrin on this loss (Figure 7B). Given that we did not observe a significant loss of ipsilateral TH^+^ neurons in females, we did not estimate NeuN counts in female mice. Consistent with TH results, we did not observe any contralateral loss of NeuN^+^ neurons in male mice (Supplementary Figure 5).

**Supplementary Figure 5:**
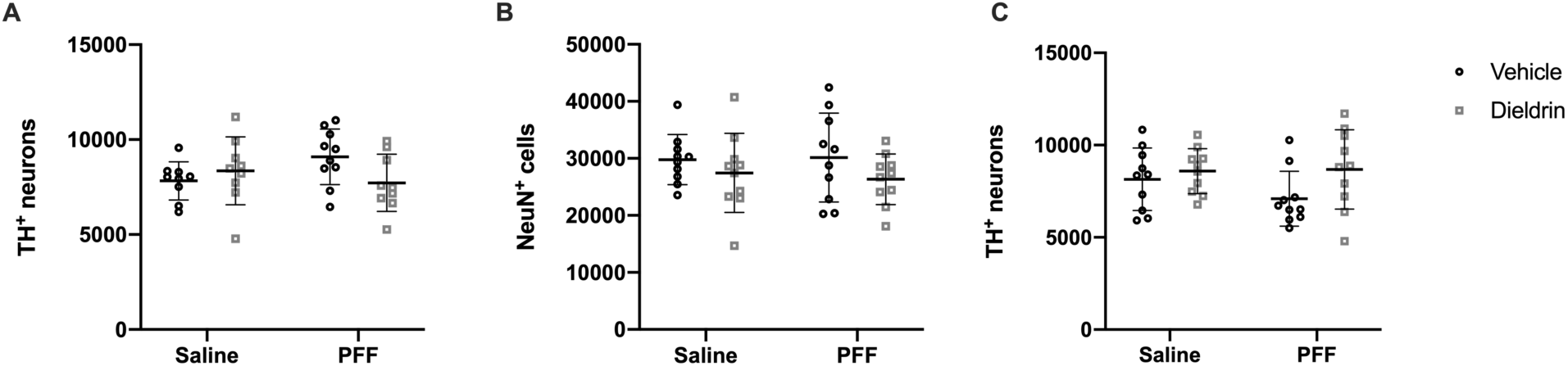
PFF injection and dieldrin show no effect on contralateral TH and NeuN immunoreactive neurons. A) Number of TH+ neurons in the contralateral nigra were estimated by unbiased stereology (n = 10 per group). A) There was no effect of dieldrin or PFF on contralateral nigral TH^+^ neurons in male animals (two-way ANOVA: PFF, p = 0.5172; dieldrin, p = 0.3922; interaction = 0.0581). B) There was no effect of dieldrin or PFF on contralateral nigral NeuN^+^ neurons in male animals two-way ANOVA: PFF, p = 0.8462; dieldrin, p = 0.1186; interaction, p = 0.7063). C) There was no effect of dieldrin or PFF on contralateral nigral TH^+^ neurons in female animals (two-way ANOVA: PFF, p = 0.3645; dieldrin, p = 0.0627; interaction = 0.2833). Data shown as mean +/- 95% CI.

### Dieldrin exposure does not alter striatal α-syn levels

Since we observed sex-specific effects of dieldrin exposure on the response to synucleinopathy, we tested whether developmental dieldrin exposure led to changes in α-syn levels in the striatum in adult male animals at 12 weeks of age (the age at which PFF injections were performed) by western blot. We did not observe an effect of developmental dieldrin exposure on levels of α-syn in the striatum of male mice developmentally exposed to dieldrin (Figure 8A,B).

**Figure 8:**
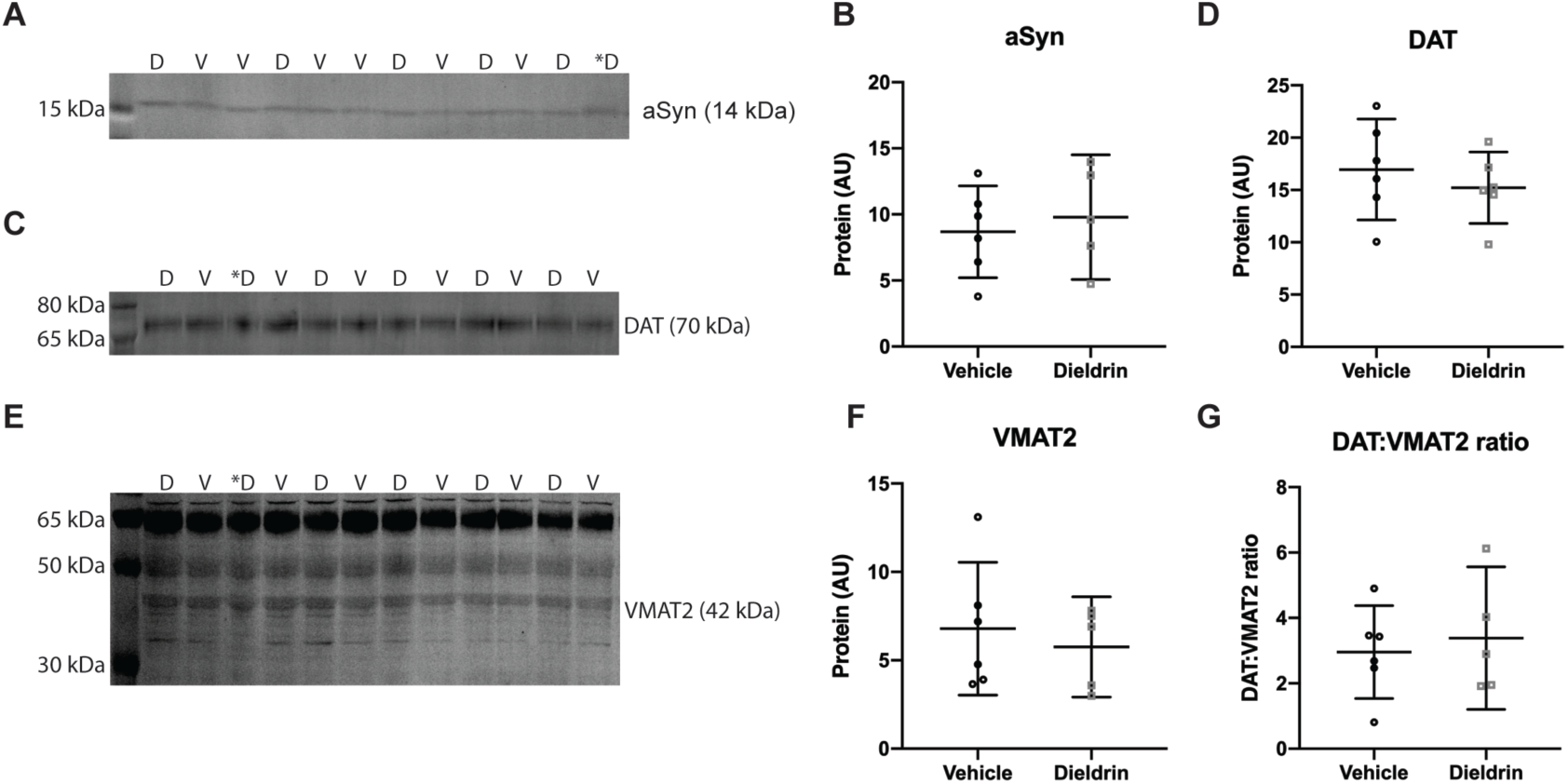
Effect of dieldrin exposure on levels of monomeric α-syn, DAT and VMAT2 in the striatum of male animals. Monomeric α-syn (A), DAT (C) and VMAT2 (E) were detected by western blot (vehicle: n = 6; dieldrin: n = 5). Samples are in mixed order for more accurate quantification. D= dieldrin, V=vehicle. Dieldrin sample with a * was excluded from all analysis. This sample was not stored properly and ran atypically on some blots. Full blots and total protein staining are shown in **Supplementary Figure 6.** B) Quantification shows no effect of dieldrin on α-syn levels in the striatum (unpaired t-test with Welch’s correction: p = 0.6279). D) Quantification shows no effect of dieldrin on DAT levels in the striatum (unpaired t-test with Welch’s correction: p = 0.8469). F) Quantification of the 42 kDa band of dieldrin shows no effect of dieldrin on VMAT2 levels in the striatum (unpaired t-test with Welch’s correction: p = 0.5764). G) Dieldrin shows no effect on DAT:VMAT2 ratio (unpaired t-test with Welch’s correction: p = 0.6700). Data shown as mean +/- 95% CI.

### Dieldrin exposure does not alter striatal DAT and VMAT2 levels

In the 2006 study showing that developmental dieldrin exposure exacerbated MPTP toxicity, the authors observed a dieldrin-induced increase in the DAT:VMAT2 ratio and a corresponding increase in DA turnover.^33^ To test if these findings were replicated in our experiment, we performed western blots for DAT and VMAT2 from striatum of male mice. In contrast to these previous results, we did not observe a dieldrin-induced change in DAT, VMAT2 or the DAT:VMAT2 ratio in our study (Figure 8C-G).

**Supplementary Figure 6:**
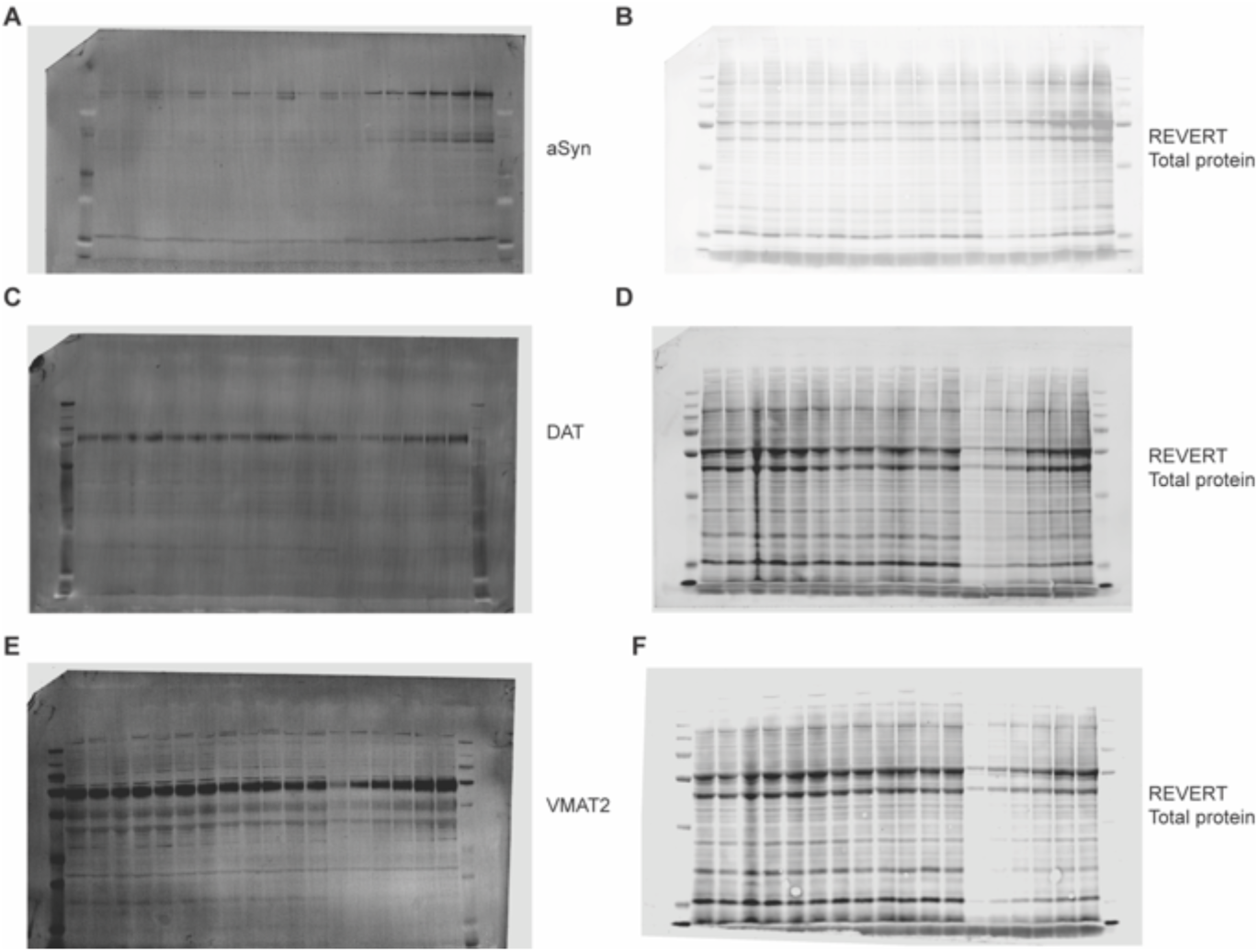
Western blots and total protein staining for Figure 8. α-synuclein (A) and corresponding REVERT total protein stain (B). DAT (C) and corresponding REVERT (D). VMAT2 (E) and corresponding REVERT (F). Co-blotted dilutions standards are also shown in the last six lanes on the right side of each blot.

### Dieldrin exposure induces sex-specific patterns of expression in inflammatory genes

There is a growing recognition that neuroinflammation plays an important role in human PD and in the α-syn PFF model.^66^ In addition, previous results demonstrated that while dieldrin exposure did not affect glial fibrillary acidic protein (GFAP) levels in the striatum, it did exacerbate MPTP-induced increases in GFAP, suggesting that dieldrin exposure leads to a greater neuroinflammatory response to a second insult.^33^ Thus, we sought to determine whether developmental dieldrin exposure affects expression of neuroinflammatory genes in the striatum. We screened the expression of a targeted set of neuroinflammatory genes in striata from male and female mice developmentally exposed to dieldrin using the TaqMan Array Card Mouse Immune Panel. Analysis was stratified by sex to assess whether dieldrin has sex-specific effects on inflammatory gene expression. We observed distinct sex-specific effects on expression of neuroinflammatory genes, consistent with our previous results reporting sex-specific effects on DNA methylation and the transcriptome in the ventral midbrain ^35^. In male mice, nine genes were differentially expressed by dieldrin exposure (p ≤ 0.05) (Table **3**). In female mice, 18 genes were differentially expressed by dieldrin exposure (Table **4**).

**Table 3:**
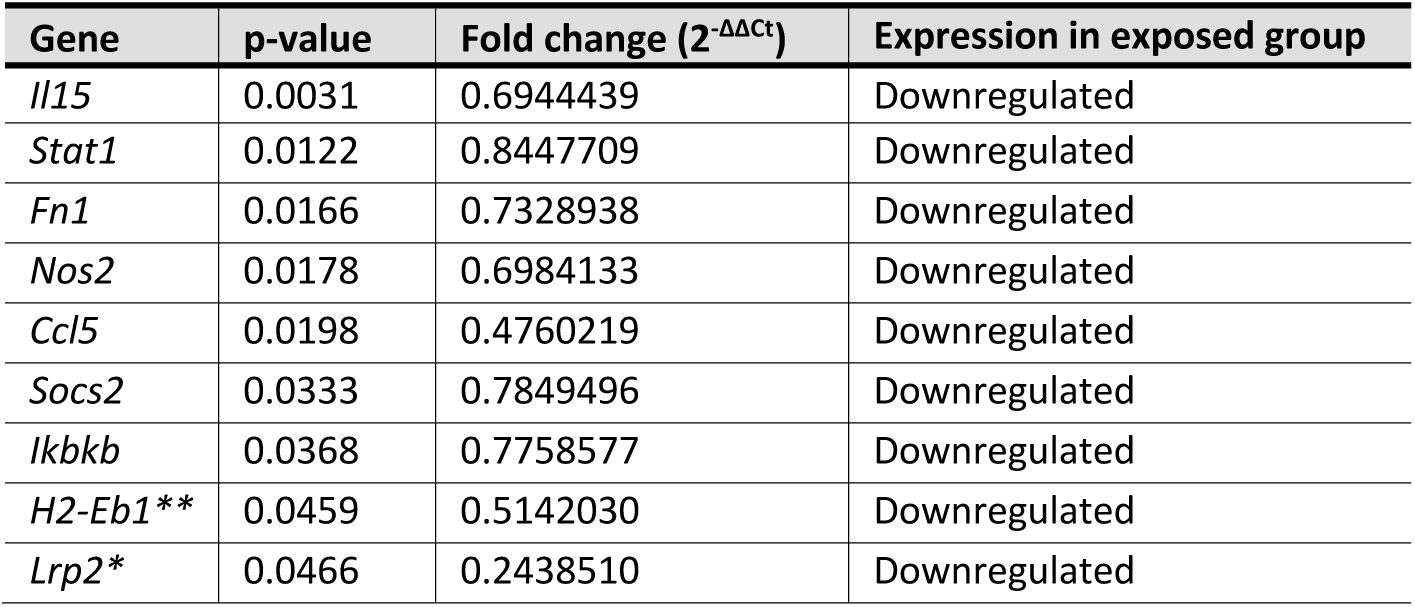
Differentially regulated genes in males. Genes that were differentially regulated between the dieldrin exposed group vs the control group in male mice (n=8 per treatment group, p < 0.05). One star (*) indicates the genes that do not cluster with other genes in STRING network analysis; two stars (**) indicates that the gene was not mapped to the STRING database.

To investigate whether the identified differentially expressed genes (DEGs) have known interactions, we performed STRING protein-protein network analysis. STRING analysis showed that 7 of the 9 (77.8%) DEGs in males have known interactions between their encoded proteins (Figure 9A). Meanwhile, 16 of the 18 (88.8%) DEGs in females have known interactions between their encoded proteins (Figure 9B). Since this was a curated group of genes selected for function, we expected this high degree of connectivity and a high number of significantly enriched gene ontology terms. For both networks, the most enriched gene ontology terms were related to the cellular response to cytokines (Supplementary Table 1,2).

**Figure 9:**
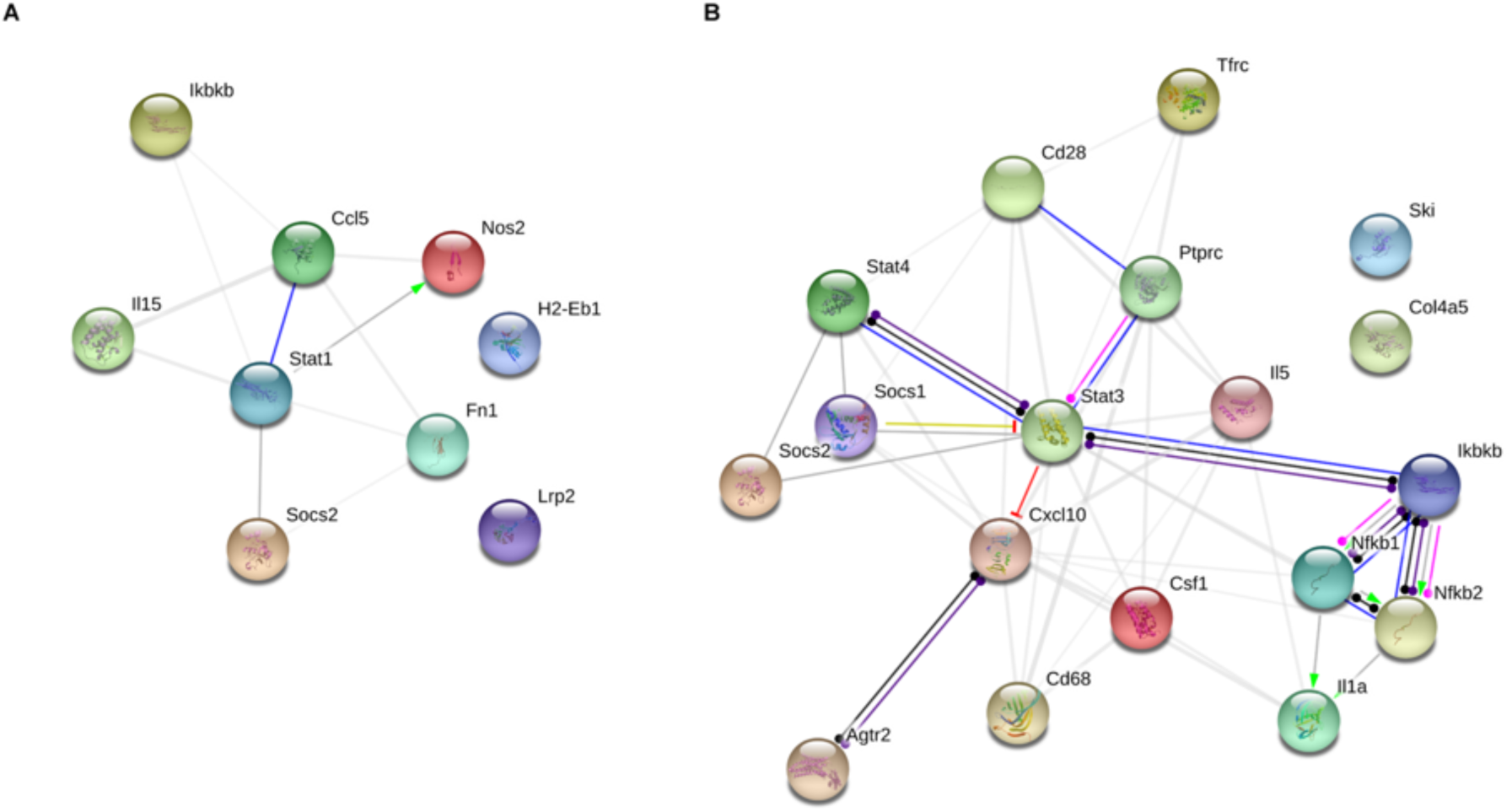
STRING interaction networks for male and female differentially expressed genes. Dieldrin-related differentially expressed genes (DEGs) for male and female animals were placed into the STRING network tool to investigate known interactions between the proteins encoded by the genes. We found a high degree of interconnectivity for both the male and female gene lists. A) In males, 7 of the 9 (77.8%) DEGs had known interactions. B) In females, 16 of the 18 (88.8%) DEGs had known interactions.

## Discussion

### Male-specific exacerbation of synucleinopathy-induced deficits in motor behavior developmental dieldrin exposure

As an important validation, our results in the PFF-injected animals without dieldrin exposure replicated previous reports of this model in mice. Specifically, we observed comparable levels and timing of PFF-induced accumulation of psyn-positive aggregates, loss of striatal DA, and loss of nigral TH^+^ cells.^38^ Our new results in dieldrin-exposed animals demonstrate that developmental dieldrin exposure induces a male-specific exacerbation of PFF-induced toxicity consistent with previous results in the MPTP model.^33^ However, dieldrin did not exacerbate the timing or extent of PFF-induced α-syn accumulation and aggregation into p-syn positive aggregates in the nigra (Figure 4), indicating that dieldrin does not affect the propensity of α-syn to aggregate, but instead affects the response to the aggregation. This exacerbated response manifests as increased PFF-induced motor deficits assessed on the challenging beam and deficits in striatal DA handling 6 months post-PFF injection in animals exposed to dieldrin in male mice (Figure 3, Figure 5).

Of note, we showed that PFF injection alone did not affect the speed of male mice on the challenging beam. Only PFF-injected animals previously exposed to dieldrin showed an effect on this outcome measure, displaying a longer time to traverse at 6 months post-PFF injection, suggesting a more severe dopaminergic deficit in PFF-injected animals previously exposed to dieldrin (Figure 3A). This finding is similar to results in other mouse models of PD-related pathology, including the MPTP mouse model and the Pitx3-aphakia mouse.^55,58^ Consistent with previous results in other α-syn models including the Thy1-α-syn overexpression models, we also found that PFF injection alone induced a significant increase in errors made on the beam (Figure 3C).^54,67^ While the PFF-injected animals previously exposed to dieldrin did make more errors at 6 months compared to baseline, indicating a worsening of sensorimotor function, they did not make as many errors compared to the PFF-injected animals not exposed to dieldrin (Figure 3C, Supplementary Figure 2C). This seeming discrepancy in results on time to traverse and errors is actually consistent with our previous observations. In MPTP-treated mice, we have observed that this increase in time to traverse can be associated with a reduced number of errors (Fleming et al. unpublished observations). MPTP-treated mice move more slowly across the length of the beam and appear to be more “cautious” with their stepping, resulting in fewer mistakes. Thus, these results support the hypothesis that dieldrin exposure exacerbates PFF-induced motor deficits. All of the observed impairments on challenging beam were specific to male mice, with female mice showing no effect of dieldrin exposure or PFF injection on their performance on challenging beam (Figure 3; see ***Sex differences in PFF-induced motor deficits*** below).

### Male-specific exacerbation of synucleinopathy-induced deficits in DA turnover after developmental dieldrin exposure

Defects in DA handling are broadly indicative of DA neuron dysfunction; consistent with this, we observed PFF-induced deficits in the DOPAC:DA and HVA:DA ratios, measures of DA turnover and handling, in both male and female mice (Figure 5D,E,I,J). None of these outcome measures were exacerbated by dieldrin exposure except for HVA:DA in male mice at 6 months after PFF injections. This male-specific exacerbation of the PFF-induced increase in DA turnover is indicative of a greater stress on the DA system in male animals and consistent with the male-specific exacerbation of motor deficits discussed above. In addition, the male-specific deficit in DA turnover is consistent with a large body of evidence supporting a central role for cytosolic DA in PD pathogenesis.^68^ Critically, it is not only overall levels of DA that matter for disease etiology and progression, but also the ability of a neuron to maintain DA within synaptic vesicles to both protect against cytosolic degradation of DA and allow the cell to release enough DA into the synapse.^69–79^ Taken together, the behavioral and HPLC data suggest that developmental dieldrin exposure causes persistent changes to the nigrostriatal system that exacerbate the response to PFF-induced synucleinopathy through disruption of striatal synaptic terminals (Figure 6, Figure 10). These results suggest that a more detailed analysis of DA uptake, release and turnover in the striatum is warranted in this two-hit model.

**Figure 10:**
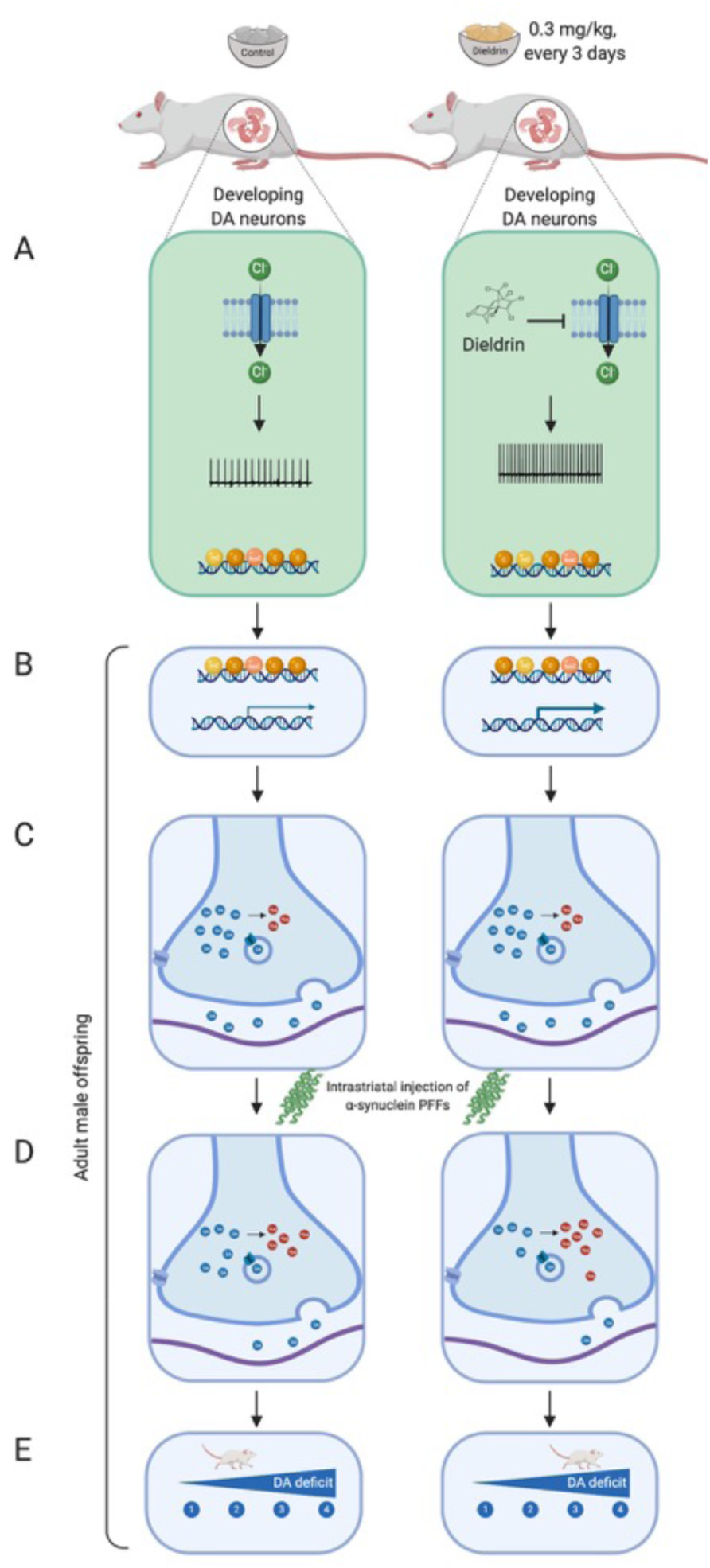
Proposed mechanism by which developmental dieldrin exposure leads to exacerbation of PFF-induced toxicity. Dams are fed vehicle or dieldrin containing food starting 1 month prior to mating and continuing through weaning of F1 pups. Dieldrin inhibits chloride influx through GABA_A_ receptors resulting in increased neuronal activity (A). This change in activity produces epigenetic changes throughout the lifespan even when dieldrin is no longer present (B). These epigenetic changes affect dopamine neuron development and maintenance, producing stable changes in striatal dopamine synapse function (C). These synaptic changes lead to increased susceptibility to PFF-induced synucleinopathy (D, E). Created in BioRender.

In contrast to the neurochemical findings, we did not observe an exacerbation in loss of nigral TH phenotype or degeneration of nigral neurons at the same 6-month time point (Figure 7). The discrepancy between these striatal and nigral observations may be a byproduct of the time point chosen. Six months after PFF injections was the latest time point that we assessed in this study, but the earliest time point where we observed neurochemical or behavioral deficits. It is possible that 6 months old is still too early to observe the effects of exposure-induced exacerbation of degeneration, which typically lags behind striatal dysfunction and degeneration of the synaptic terminals. Thus, at later time points, animals with greater striatal synaptic deficits may eventually show greater nigral loss. Now that we have established the phenotype in this two-hit model, further studies will assess possible acceleration and/or exacerbation of these synaptic deficits in the striatum at later time points.

### A model of dieldrin-induced increases in neuronal susceptibility

Based on the results reported here, our previous characterization of epigenetic changes induced by developmental dieldrin exposure, and the mechanisms of dieldrin toxicity, we propose a model for how developmental dieldrin exposure leads to increased susceptibility to synucleinopathy.^33,35^ In this model, exposure to dieldrin occurs during prenatal and postnatal development. The half-life of dieldrin in mouse brain is less than a week, so no detectable dieldrin remains in the brain of F1 offspring by a few weeks after weaning.^33,34,80^ When dieldrin is present in the developing brain, it is thought to act on developing DA neurons by inhibiting GABA_A_ receptor-mediated chloride flux, resulting in increased neuronal activity (Figure 10A).81– ^86^ Based on previous results, we propose that this net increase in neuronal activity modifies the dopamine system through persistent changes in epigenetic mechanisms, leading to dysregulation of genes important for dopamine neuron development and maintenance (Figure 10B,C).^35^ These stable changes then alter the susceptibility of this system to future insults, likely via alterations in striatal dopamine synapses that manifest as increased DA turnover upon application of PFFs (Figure 5, Figure 10C). Supporting this model, the present work identified a dieldrin-induced, male-specific exacerbation of PFF-induced deficits in striatal DA handling (Figure 5, Figure 10D) and motor behavior (Figure 3, Figure 10E). Further studies in our lab will focus on exploring the synaptic mechanisms underlying this phenotype and will aim to connect the observed epigenetic changes with these mechanisms. In addition, in a current study, we are tracking the longitudinal patterns of dieldrin-induced epigenetic changes from birth to 12 weeks of age to determine if dieldrin-induced epigenetic changes are maintained from birth or if they represent an altered longitudinal trajectory of epigenetic changes.

### Dieldrin induced sex-specific effects in the nigrostriatal pathway may underlie the male-specific increase in susceptibility

As discussed above, work in our lab has focused on characterizing dieldrin-induced changes in the DA system that may underlie this increase in susceptibility.^35^ In a previous study, we reported sex-specific, dieldrin-associated changes in DNA methylation and gene transcription in the ventral midbrain at genes related to dopamine neuron development and maintenance. These dieldrin-induced changes in gene regulation were identified at 12 weeks of age, which is when male-specific exacerbation of PFF- and MPTP-induced toxicity is observed.^33^ To complement those results, we explored additional dieldrin-induced changes in littermates in our study that did not receive PFF or saline injections.

Consistent with our finding that dieldrin did not exacerbate PFF-induced α-syn aggregation, we also did not observe dieldrin-induced changes in overall levels of α-syn (Figure 8). This finding replicates previous results in this exposure paradigm.^33^ In contrast to the findings in Richardson et al, we did not observe dieldrin-induced changes in overall levels of DAT or VMAT2, or in the DAT:VMAT2 ratio in male animals (Figure 8). The previous report showed increased DAT and VMAT2 in both male and female animals exposed to the same dose of dieldrin used here, as well as a male-specific increase in DAT:VMAT2 ratio.^33^ Despite this difference in results, we observed a similar male-specific exacerbation of toxicity upon application of the second hit (PFFs in this study, MPTP in the previous study).

Next, we tested expression of a curated set of inflammatory genes in the striatum of dieldrin-exposed animals to determine if developmental dieldrin exposure caused long lasting changes in the neuroinflammatory system in adulthood. Recognition of an important role of neuroinflammation in human PD and in the α-syn PFF model has been growing.^66,87–95^ In addition, previous results demonstrated that dieldrin exposure exacerbates MPTP-induced increases in expression of glial fibrillary acidic protein (GFAP) levels in the striatum, suggesting that dieldrin exposure leads to a greater neuroinflammatory response to a second insult. While we were unable to test expression of neuroinflammatory genes after the application of PFFs, we did identify dieldrin-induced changes in the expression of neuroinflammatory genes. Consistent with our previous results showing sex-specific effects of dieldrin exposure on the nigral epigenome and transcriptome, we identified sex-specific effects of dieldrin on neuroinflammatory gene expression (n=9 in male mice; n=18 in female mice) (Table 3, Table 4). Since this was a curated set of genes, we also observed a very high degree of connectivity between these genes in STRING protein-protein network analysis (77.8% of male DEGs and 88.8% of female DEGs) (Figure 9). For both networks, the most enriched gene ontology terms were related to the cellular response to cytokines (Supplementary Table 1,2).

**Table 4:** Differentially regulated genes in females. Genes that were differentially regulated between the dieldrin exposed group vs the control group in female mice (n=8 per treatment group, p < 0.05). One star (*) indicates the genes that do not cluster with other genes in STRING network analysis; two stars (**) indicates that the gene was not mapped to the STRING database.

| Gene | p-value | Fold change ($2^{-\Delta\Delta Ct}$) | Expression in exposed group |
| --- | --- | --- | --- |
| *Csf1* | 0.0003 | 1.5955447 | Upregulated |
| *Tfrc* | 0.0024 | 1.2815964 | Upregulated |
| *Agtr2* | 0.0054 | 1.9362228 | Upregulated |
| *Stat4* | 0.0055 | 1.7925272 | Upregulated |
| *Cd68* | 0.0060 | 0.6350103 | Downregulated |
| *Socs1* | 0.0075 | 0.7602232 | Downregulated |
| *Ptprc* | 0.0078 | 1.3596591 | Upregulated |
| *Ikbkb* | 0.0140 | 0.7839894 | Downregulated |
| *Nfkb2* | 0.0198 | 1.2670066 | Upregulated |
| *Col4a5\*\** | 0.0222 | 1.4464285 | Upregulated |
| *Nfkb1* | 0.0238 | 1.2724289 | Upregulated |
| *Cxcl10* | 0.0337 | 2.0286149 | Upregulated |
| *Il1a* | 0.0351 | 0.8374558 | Downregulated |
| *Cd28* | 0.0382 | 0.4077085 | Downregulated |
| *Stat3* | 0.0419 | 1.1916636 | Upregulated |
| *Il5* | 0.0437 | 2.6215317 | Upregulated |
| *Socs2* | 0.0465 | 1.2408090 | Upregulated |
| *Ski\** | 0.0482 | 1.2297090 | Upregulated |

No single inflammatory pathway is apparent in the list of differentially regulated genes from either sex and these results are not consistent with canonical pro- or anti-inflammatory effects. In interpreting these results, it is critical to remember that gene expression was measured in developmentally exposed offspring at 12 weeks of age, when dieldrin is no longer detectable in the brain. Thus, these expression changes may not reflect a typical acute or even chronic inflammatory response. As with our previous epigenetic study, these observed changes likely reflect a persistent change in the baseline state of this system, such that the system responds differently to the second hit.

Despite the lack of a clear pro- or anti-inflammatory gene signature, a few patterns can be identified in the list of DEGs. Only one gene is differentially expressed in both sexes – *IKBKB*, which encodes an NF-κB inhibitor. This gene is downregulated in both male and female animals, but only female animals show a corresponding increase in *NKFB1* expression. The combination of increased *NKFB1* expression and decreased *IKBKB* in the female animals suggests a state of microglial activation. Consistent with this idea, we observed increased expression of the microglial pro-inflammatory cytokine genes, *CXCL10* and *CSF1*. However, we also observed dieldrin-induced decreased expression of *IL1A* and increased expression of *SOCS1*, changes that are not consistent with a pro-inflammatory state. In males, the downregulation of *IKBKB* is not accompanied by corresponding pro-inflammatory changes. Instead, we observed decreased expression of four pro-inflammatory genes – *IL15, STAT1, NOS2*, and *CCL5.* We also observed expression changes in genes involved in the adaptive immune response (upregulation of *IL5, PTPRC, STAT3*, and *STAT4*, and downregulation of *CD28*) in female animals. When considered together, these gene expression results establish that developmental dieldrin exposure induces distinct sex-specific effects on neuroinflammatory pathways. While these changes are not consistent with canonical pro- or anti-inflammatory effects, they provide multiple avenues for follow-up studies. In particular, future studies will determine if these observed changes in gene expression correspond with activation or deactivation of microglia or the adaptive immune system in dieldrin-exposed animals. Furthermore, using our two-hit model, follow-up studies will test whether specific dieldrin-induced DEGs respond differently to a PFF second hit or if dieldrin exposure modifies PFF-induced microglial activation or immune response.

### Sex differences in PFF-induced motor deficits

While we expected to see a male-specific exacerbation of PFF-induced toxicity by developmental dieldrin exposure, the finding that PFFs alone did not induce motor deficits in female animals was quite surprising. Indeed, this is the first study to show sex differences in sensorimotor function in the PFF model. Despite male and female animals showing similar levels of PFF-induced psyn aggregation (Figure 4), loss of DA, DOPAC and HVA levels and increases in DA turnover (Figure 5, Supplementary Figure 3), female mice showed no motor behavior deficits (Figure 3). While these data at first appear contradictory, we found that female animals showed a ∼20% loss of ipsilateral nigral TH immunoreactivity between saline- and PFF-injected groups, whereas males showed a ∼35% PFF-induced loss of TH immunoreactivity, ipsilateral to PFF injection (Figure 7). Together, these data suggest that female mice have reduced or slower loss of nigral DA neurons and a behavioral resilience to the same level of DA loss and similar defects in DA turnover compared to their male counterparts. This is consistent with the reduced incidence of PD and severity of disease course in human females.^96–105^ Collecting tissue at a later time point (e.g. 9 months post-PFF injection) would reveal if the female-specific resilience is complete or simply reflects a slower progression of the effects of the observed neuropathology. This unanticipated finding is particularly important as it suggests that the PFF model may be a valuable tool to model sex-differences in PD pathology and etiology that does not require any additional surgery or other treatments to manipulate hormonal state. Although this study was not designed to directly compare male and female animals, these results warrant further investigation into sex differences in the PFF model.

## Conclusions

In this paper, we demonstrated sex-specific effects of developmental dieldrin exposure on α-syn PFF-induced toxicity. Specifically, we showed that developmental dieldrin exposure increases α-syn-PFF-induced motor deficits and deficits in DA handling but does not affect PFF-induced loss of nigral TH^+^ neurons. These results indicate that our two-hit exposure model represents a novel experimental paradigm for studying how environmental factors increase risk of PD. In addition, we observed sex-specific effects of developmental dieldrin exposure on neuroinflammation, and a female-specific resilience to PFF-induced pathology. These sex-specific effects are likely not specific to the toxicants used here. A recent study demonstrated similar sex differences in response to rotenone, a commonly studied parkinsonian toxicant.^106^ Given the reduced incidence of PD and severity of disease course in human females, the sex differences in these models underscore the need to include female animals in toxicity studies.^96–105^

## Author Contributions

AIB planned and designed all experiments. AOG and SEV carried out all mouse husbandry and dieldrin dosing, and JK assisted. PFF surgeries were planned by CES and performed by AOG, AB, CES, JRP, and KMM, with surgical assistance by JK, SEV and CJK. KCL provided PFFs. AOG performed all experimental outcomes except for those listed here. NCK performed NeuN stereology. DEH, AM and SMF scored and analyzed motor behavior. ACS and JWL performed HPLC analysis. ACS isolated RNA and did qPCR assays. JK analyzed qPCR data. AIB and AOG wrote the manuscript. JK, CES, JWL, KCL, SMF, JPR, CJK edited and provided feedback on the manuscript

## Funding

This work was supported by the National Institutes of Health (R21 ES029205, R00 ES024570, R33 NS099416).

## Supplemental Materials

Supplementary table 1: String gene ontology results for male differentially expressed genes Supplementary table 2: String gene ontology results for female differentially expressed genes This project was pre-registered at Open Science Framework (<u>10.17605/OSF.IO/H49XF</u> and <u>10.17605/OSF.IO/WPF6U</u>).

All data for published figures and supplementary figures is deposited in Mendeley Data. GraphPad Prism files can be viewed in the free Viewer mode or data can be extracted by viewing files in a text editor. R and RStudio are freely available.

## Notes

#### Summary of Updates

Corrected author info, modified Figure 8 with correct graphs

https://doi.org/10.17605/OSF.IO/WPF6U

https://doi.org/10.17605/OSF.IO/H49XF

